# SiFit: A Method for Inferring Tumor Trees from Single-Cell Sequencing Data under Finite-site Models

**DOI:** 10.1101/091595

**Authors:** Hamim Zafar, Anthony Tzen, Nicholas Navin, Ken Chen, Luay Nakhleh

## Abstract

Single-cell sequencing (SCS) enables the inference of tumor phylogenies that provide insights on intra-tumor heterogeneity and evolutionary trajectories. Recently introduced methods perform this task under the infinite-sites assumption, violations of which, due to chromosomal deletions and loss of heterozygosity, necessitate the development of inference methods that utilize finite-site models. We propose a statistical inference method for tumor phylogenies from noisy SCS data under a finite-sites model. The performance of our method on synthetic and experimental datasets from two colorectal cancer patients to trace evolutionary lineages in primary and metastatic tumors suggest that employing a finite-sites model leads to improved inference of tumor phylogenies.

## Background

Intra-tumor heterogeneity, as caused by a combination of mutation and selection [1–4], poses significant challenges to the diagnosis and clinical therapy of cancer [5–8]. This heterogeneity can be readily elucidated and understood if the evolutionary history of the tumor cells was known. This knowledge, alas, is not available, since genomic data is most often collected from one snapshot during the evolution of the tumor’s constituent cells. Consequently, using computational methods that reconstruct the tumor phylogeny from sequence data is the approach of choice. However, while intra-tumor heterogeneity has been widely studied, the inference of a tumor’s evolutionary history remains a daunting task.

Most studies to-date relied on bulk high-throughput sequencing data, which represents DNA extracted from a tissue consisting millions of cells [9–13]. As a result, the admixture signal obtained from such data represents an average of all the distinct subpopulations present in the tumor [14]. This ambiguity makes it difficult to identify the lineage of the tumor from the mixture. In such cases, phylogenetic reconstruction requires a deconvolution of the admixture signal to identify the taxa of the tree [15–17]. This type of data is low-resolution and can not depict cellto-cell variability that is needed for inference of tumor evolution [14, 18]. Another approach for resolving intra-tumor heterogeneity and reconstructing tumor phylogeny is multi-region sequencing, in which, DNA sampled from multiple spatially separated regions of the tumor are sequenced [19, 20], however, this approach is restricted to cases where the subpopulations are geographically segregated and can not resolve spatially intermixed heterogeneity [21].

### Single-cell DNA sequencing: promises and challenges

With the advent of single-cell DNA sequencing (SCS) technologies, high-resolution data are becoming available, which promise to resolve intra-tumor heterogeneity to a single-cell level [14, 18, 22–25]. These technologies provide sequencing data from single cells, thus allowing for the reconstruction of the cell lineage tree. However, high error rates associated with single-cell sequencing data significantly complicates this task.

The whole-genome amplification (WGA) process, a crucial step in producing single-cell sequencing data, introduces different types of noises that result in erroneous genotype inferences. The prominent WGA errors include: allelic dropout (ADO) errors, false positive errors (FPs), non-uniform coverage distribution and low coverage regions [14]. Allelic dropout is a prominent error in SCS data, which contributes a considerable amount of false negatives in point mutation datasets. ADO is responsible for falsely representing the heterozygous genotypes as homozygous ones and the extent of such errors varies from 0.0972 to 0.43 as reported in different SCS-based studies [22–26]. Even though variant callers have been proposed for reducing ADO errors [27], the extent of such errors is still large. Different single-cell sequencing studies have reported false positive rates varying from 1.2 × 10^-6^ to 6.7 × 10^-5^ [22–26], the number of occurrences of which can essentially exceed the number of true somatic mutations. Often consensus-based approach is taken to reduce the number of false positive errors [26–28], in which, variants only observed in more than one single cell are considered. The variants observed in only one single cell are treated as errors and removed. In doing so, this approach also removes the true biological variants unique to a cell whereas, sites of recurrent errors persist. Both ADO and coverage non-uniformity result in unobserved sites. Often more than 50% of the genotypes are reported as missing due to the low quality of SCS data and thus no information regarding the mutation status of that site is conveyed [22]. Another source of error in SCS data are ‘cell doublets’ in which two or more cells are accidentally isolated instead of single cells. The cell doublet error rates vary considerably depending on the isolation technology. Methods such as FACS have reported less than 1% cell doublet error rates [29–31], while doublet rates for methods such as mouth pipetting and microdroplet encapsulation technologies range from 1-10% [22, 23, 32].

### Existing work

Single-cell-based studies for delineating the tumor phylogeny rely on the single-cell somatic SNV profiles, which are confounded by the technical errors in single-cell sequencing. Even though such errors prohibit the use of classic phylogenetic approaches, many studies have used them. Distance-based methods like UPGMA and neighbor joining have been used by Yu *et al.* [33], and Xu *et al.* [23] respectively. Eirew *et al.* [34] used a popular Bayesian phylogenetic inference tool, MrBayes [35], for inferring evolutionary history. However, none of these methods account for the SCS specific errors.

BitPhylogeny [36] is a non-parametric Bayesian approach that uses a tree-structured mixture model to infer intra-tumor phylogeny. Even though such an approach is valuable for identifying subclones from bulk sequencing data, it is not suitable in the context of present-day single-cell datasets (fewer than 100 cells) [22–24, 26, 33], which do not provide sufficient data required by the mixture model in order to converge to the target distribution [37]. Furthermore, BitPhylogeny is a flexible framework that can fit different data types but does not specifically model the single-cell errors.

SCITE [38] and OncoNEM [37] are two computational tools that were specifically designed for inference of tumor evolution from SCS data. SCITE is an MCMC algorithm that allows one to infer maximum likelihood tree from imperfect genotype matrix of SCS. It infers the evolutionary history as a mutation tree, proposed by Kim and Simon in [39]. A mutation tree shows the chronological order of the mutations that occur during tumor development. OncoNEM is a likelihood-based method that employs a heuristic search algorithm to find the maximum likelihood clonal tree, a condensed tree that represents the evolutionary relationship between the subpopulations in the data. OncoNEM clusters the cells together into clones and also infers unobserved populations that can improve the likelihood. Both of these methods probabilistically account for technical errors in SCS data and can also estimate the error rates of SCS data. However, both SCITE and OncoNEM suffer by making inferences under the “infinite sites assumption”, which posits that each site in the dataset mutates at most once during the evolutionary history [40] and the taxa form a perfect phylogeny [41]. This assumption is often violated in human tumors due to different events such as chromosomal deletions, loss of heterozygosity (LOH) and convergent evolution [42]. Furthermore, OncoNEM infers clonal trees where cell-to-cell evolution is not displayed, and SCITE is concerned with the order of mutation in the tree but not the lineage of single cells. To the best of our knowledge, there is no method that infers a phylogenetic tree from SCS data under finite-site model of evolution while accounting for the technical errors in SCS.

### SiFit

Here we propose SiFit, a likelihood-based approach for inferring tumor trees from imperfect SCS genotype data with potentially missing entries, under finite-site model of evolution. To account for the errors in SCS, SiFit extends the error model of SCITE and OncoNEM. This extension accommodates for the possible genotypes that are excluded by infinite sites model. SiFit employs a finite-site model of evolution that accounts for the effects of deletion, LOH and point mutations on the genomic sites via transition probabilities between genotype states. SiFit employs a heuristic search algorithm to find the phylogenetic tree that is most likely to produce the observed SCS data. We evaluate SiFit on a comprehensive set of simulated data, where it performs superior to the existing methods in terms of tree reconstruction. Application of SiFit to experimental datasets shows how infinite sites assumption gets violated in real SCS data and how SiFit’s reconstructed tumor phylogenies are more comprehensive compared to phylogenies reconstructed under infinite sites assumption. SiFit achieves a major advance in understanding tumor phylogenies from single cells and is applicable to wide variety of available single-cell DNA sequencing datasets.

## Results and discussion

### Overview of SiFit

We start with a brief explanation of how SiFit infers a tumor phylogeny from noisy genotype data obtained from single-cell sequencing. The input data consist of the following: (1) an *n × m* genotype matrix, which contains the observed genotypes for *m* single cells at *n* different loci, the genotype matrix can be binary or ternary depending on the data, and (2) the false positive rate (FPR), *α* and false negative rate (FNR), *β*. These error parameters can be learned from the data.

SiFit includes (1) a finite-site model of tumor evolution and an error model for SCS, based on which the likelihood score of a candidate phylogenetic tree and error rate can be quantified and (2) a heuristic algorithm for exploring the joint space of trees and error rates in search of optimal parameters.

SiFit outputs a phylogenetic tree describing the evolutionary relationship between the single cells and the estimated error rates. The single cells are placed at the leaves of the phylogenetic tree. A more detailed technical description of SiFit can be found in the “Methods” section.

### Phylogenetic trees and model of tumor evolution

We assume that the observed single cells evolved according to an underlying phylogenetic tree. A phylogeny or phylogenetic tree represents the genealogical relationship among genes, species, populations, etc. [43]. In the context of tumor, it is a rooted binary tree that represents the genealogical relationship among a set of cells. The sequenced single cells are placed at the leaves of the phylogenetic tree. We also assume that the cells evolve according to a finite-site model along the branches of the tree.

The *n × m* true genotype matrix *G* contains the true genotypes of *m* single cells at *n* different loci. If the data contains information only about the presence or absence of a mutation at a locus, the matrix is binary, where the absence or presence of a mutation is represented by a 0 or 1 at the entry *G*(*i, j*), respectively. Assuming the cells to be diploid, if the data differentiates between heterozygous and homozygous mutations, the genotype matrix is ternary, where a 0, 1 or 2 at entry *G*(*i, j*) denotes homozygous reference, heterozygous or homozygous non-reference genotype, respectively. Heterozygous or homozygous non-reference genotypes represent mutations. This ternary representation facilitates the use of mutation profile from modern variant calling algorithms (e.g., Monovar [27] and GATK [44]) that report mutation status of a sample in terms of genotypes.

To accommodate SCS data, we develop a finite-site model of evolution (*ℳ*) that accounts for the effects of point mutations, deletion and LOH on genomic sites. The finite-site model of evolution encompasses a continuous-time Markov chain that assigns a transition probability for one genotype state changing to another along a branch of length *t*. The value of the transition probabilities depends on the branch length (*t*) and the parameters, *ℳ*_*λ*_, of the model of evolution (see “Methods” section for details). By assigning a finite probability for all possible genotype transitions, this finite-site model of evolution enables us to account for convergent evolution or reversal of genotypes that are excluded by methods that make the “infinite sites assumption” (SCITE and OncoNEM). OncoNEM also assumes only binary data and does not differentiate between heterozygous and homozygous mutations. This binarization of data might result in loss of information for a dataset with ternary genotypes as heterozygous and homozygous non-reference genotypes can not be distinguished when data is binarized. On the other hand, SCITE assumes that the observation of a homozygous non-reference genotype is due to technical errors only. These assumptions follow from using the infinite sites model and are not made by SiFit.

SCITE also removes the mutations that are present in all cells or in one cell as non-informative in tree reconstruction. SiFit does not remove such mutations as these can be informative in the computation of the likelihood under a finite-site model of evolution.

### Model of single-cell errors

The observed genotype matrix, denoted by *D*, is an imperfect noisy version of the true genotype matrix *G*. The false positive errors and the false negative errors are responsible for adding noise in the observed genotype matrix. Considering binary genotype data, false positive errors result in observing a 1 with probability *α* when the true genotype is 0. Similarly, due to false negative errors, with probability *β*, we will observe a 0, instead of a 1. These relationships between true and observed genotype matrix are given by

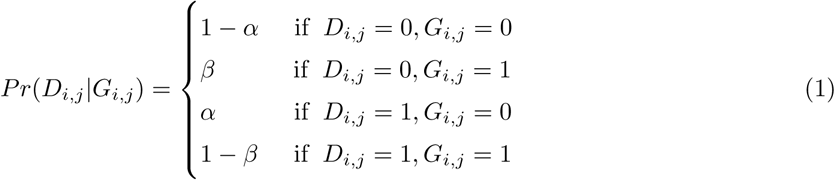

The error model for ternary data is described in detail in the “Methods” section. The observed genotype matrix can also have missing data because of uneven coverage of single-cell sequencing. SiFit handles missing data by marginalizing over possible genotypes (see “Methods” section for details).

### Tree likelihood

A phylogenetic tree, *𝒥* = (*T*, **t**) consists of a tree topology *T* and a vector of the branch lengths **t**. Assuming the technical errors to be independent of each other, and sites to evolve independently, the likelihood of a phylogenetic tree *𝒥*, the error rates ***θ*** = (*α, β*) and the parameters of the model of evolution *ℳ*_*λ*_ is given by

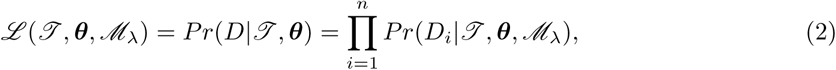

where *D*_*i*_ is the observed data at site *i* and it is a vector with *m* values corresponding to *m* single cells. The likelihood calculation for a particular site is described in detail in the “Methods” section. The maximum likelihood estimate is obtained by

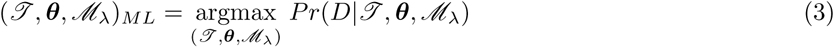

### Heuristic search algorithm

Our model has three main components, the phylogenetic tree *𝒥*, the error rates of single-cell data ***θ*** and the parameters of the model of evolution (*ℳ*_*λ*_). The tree search space has 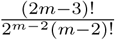 discrete bifurcating tree topologies for *m* cells, and each topology has a continuous component for branch lengths. The overall search space also has a continuous component for error rates and model parameters along with the tree space. We designed a heuristic search algorithm to explore the joint search space to infer the maximum likelihood configuration of phylogeny and error rates. In the joint (*𝒥*, ***θ****, ℳ*_*λ*_) space, we consider three types of moves to propose a new configuration. In each type of move, one component is changed. Thus from a current configuration (*𝒥*, ***θ****, ℳ*_*λ*_), a new configuration of either (*𝒥’*, ***θ****, ℳ*_*λ*_) or (*𝒥*, ***θ****’, ℳ*_*λ*_) or (*𝒥*, ***θ****, M*_*λ’*_) is proposed. The new configuration is heuristically accepted according to a ratio of likelihood. The search procedure terminates when the likelihood does not improve or the maximum number of iterations has been reached.

#### Performance on simulated data

First, we evaluated the performance of SiFit on extensive simulated datasets. The simulation studies were aimed at analyzing SiFit’s accuracy of phylogeny inference under different experimental conditions. We also assessed SiFit’s ability to estimate the error rates and its robustness against increased error rates. We compared SiFit’s performance to three other methods. To analyze how tree inference process degrades if the inference algorithm fails to account for the SCS errors, we chose a representative of classic phylogeny inference method as used by Eirew *et al.* [34]. Eirew *et al.* used MrBayes [35], a Bayesian phylogenetic inference method, that reports a set of trees drawn from the posterior distribution. Even though it was applied on SCS data, this method does not account for the errors in SCS data. The trees inferred from this method can be directly compared against the true trees. For MrBayes, we compute the average tree reconstruction error by averaging over all inferred trees. We also compared against SCITE [38] and OncoNEM [37], methods that infer tumor trees under “infinite sites assumption”. SCITE was designed to infer a mutation tree, but it can also infer a binary leaf-labelled tree, where the cells are the leaf labels and edges contain mutations. We used SCITE to infer the binary leaf-labelled tree from simulated datasets so that they can be directly compared against the true trees. Since, SCITE is an MCMC-based algorithm, occasionally it might report more than one optimal trees. In such cases, we measure average accuracy over all the reported trees. OncoNEM infers a clonal tree which can not be directly compared against the simulated trees. OncoNEM first infers a cell lineage tree and then converts it to a clonal tree by clustering nodes. The cell lineage tree inferred by OncoNEM is a different representation of the clonal tree. We convert the cell lineage tree inferred from OncoNEM to an equivalent phylogenetic tree (potentially non-binary) by projecting the internal nodes to leaves (for details see “Methods”) enabling us to compare OncoNEM results against true trees.

As the performance metric, we use the tree reconstruction error, which measures the distance of the inferred tree from the true tree. The distance between two binary trees is measured in terms of Robinson-Foulds (RF) distance [45], which counts the number of non-trivial bipartitions that are present in the inferred or the true tree but not in both the trees. We normalize this count using the total number of bipartitions in the two trees. The output of SiFit, SCITE and Bayesian phylogenetic inference algorithm (MrBayes) is compared against the true tree in terms of the RF distance. The tree inferred by OncoNEM might be non-binary, so for OncoNEM trees, we separately computed FP and FN distances between the true tree and the inferred tree. For binary trees with the same leaf set, the FP and FN distances are equal. For non-binary tree, FP and FN distances could differ from each other. The “Methods” section gives the details of the tree reconstruction error metric for comparing trees.

### Accuracy of phylogeny inference

To analyze the accuracy of SiFit’s tree inference, we simulated three sets of single-cell data with varying levels of doublet noise: (1) datasets without any doublet (*δ* = 0); (2) datasets with 5% doublet rate (*δ* = 0.05); and (3) datasets with 10% doublet rate (*δ* = 0.1). For each setting, we simulated random binary phylogenetic trees for varying number of leaves (single cells). The number of cells, i.e, leaves in the trees, *m*, was varied as *m* = 50, *m* = 100 and *m* = 200. The number of sites, *n*, was varied as *n* = 200, *n* = 400 and *n* = 600 respectively. For each combination of *δ*, *n* and *m*, we generated 10 datasets that were simulated from 10 random trees. At the root of the tree, all sites have homozygous reference genotype. The sequences are evolved along the branches of the tree starting from the root. In each branch of the tree, we simulate four types of events that can alter the genotype of a site: new mutation, deletion, loss of heterozygosity (LOH) and recurrent point mutation (see “Methods” for details). After evolving, the leaves have genotype sequences with true mutations. *m* genotype sequences corresponding to *m* single cells constitute the true genotype matrix. Errors are introduced into the true genotype matrix to simulate single-cell errors. For datasets with doublets, doublets are formed by merging the genotypes of two single cells (“Methods”) with probability *δ*. The false negative rate for cell *c*, *β*_*c*_, is sampled from a normal distribution with mean *β*_*mean*_ = 0.2 and standard deviation 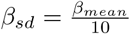. False negatives are introduced in the genotype matrix with probability *β*_*c*_ for cell *c*. We introduced false positives to the genotype matrix with error rate, *α* = 0.01, by converting homozygous reference genotypes to heterozygous genotypes with probability *α*. It is important to note that false positive rate, *α* here is by definition different from the false discovery rate (FDR) reported in single-cell-based studies such as [22, 24, 26]. *α* here indicates the fraction of non-mutant sites that are reported as mutant in the observed genotype matrix, whereas FDR reported in the aforementioned studies refers to the number of false-positive errors per sequenced base pair. For exome-sequencing studies, even a very small FDR (~ 10^-5^) can lead to a large number of false-positive variants in the observed genotype matrix making *α* much higher than the reported FDR. After adding noise, the imperfect genotype matrices were used as input to SiFit for learning maximum likelihood tree.

SiFit’s tree inference accuracy was compared against three other methods. Same imperfect genotype matrix was used as input to SiFit and SCITE. For OncoNEM and MrBayes, the genotype matrices were binarized by converting the heterozygous and homozygous non-reference genotypes to 1, i.e., presence of mutation. The comparison is shown in Fig. 1, which shows the tree reconstruction error. For each value of *n*, the mean error metric over 10 datasets is plotted along with the standard deviation as the error bar. For datasets without doublets, SiFit substantially outperforms the other three methods for all values of *m* and *n*. The performance of each algorithm except for OncoNEM improves as the value of *n* increases. The behavior of OncoNEM is different. For *m* = 100, its accuracy decreases for *n* = 600 compared to *n* = 400. This might be because, OncoNEM was developed for clonal tree inference and the effect of an additional number of sites cannot be observed in the equivalent phylogenetic tree unless they (the additional sites) are different across the clones. For datasets with higher number of sites (*n* = 600), SiFit was able to find either the true tree topology or near-perfect tree topology for most of the datasets demonstrating its ability to infer correct trees given enough data.

**Figure 1.**
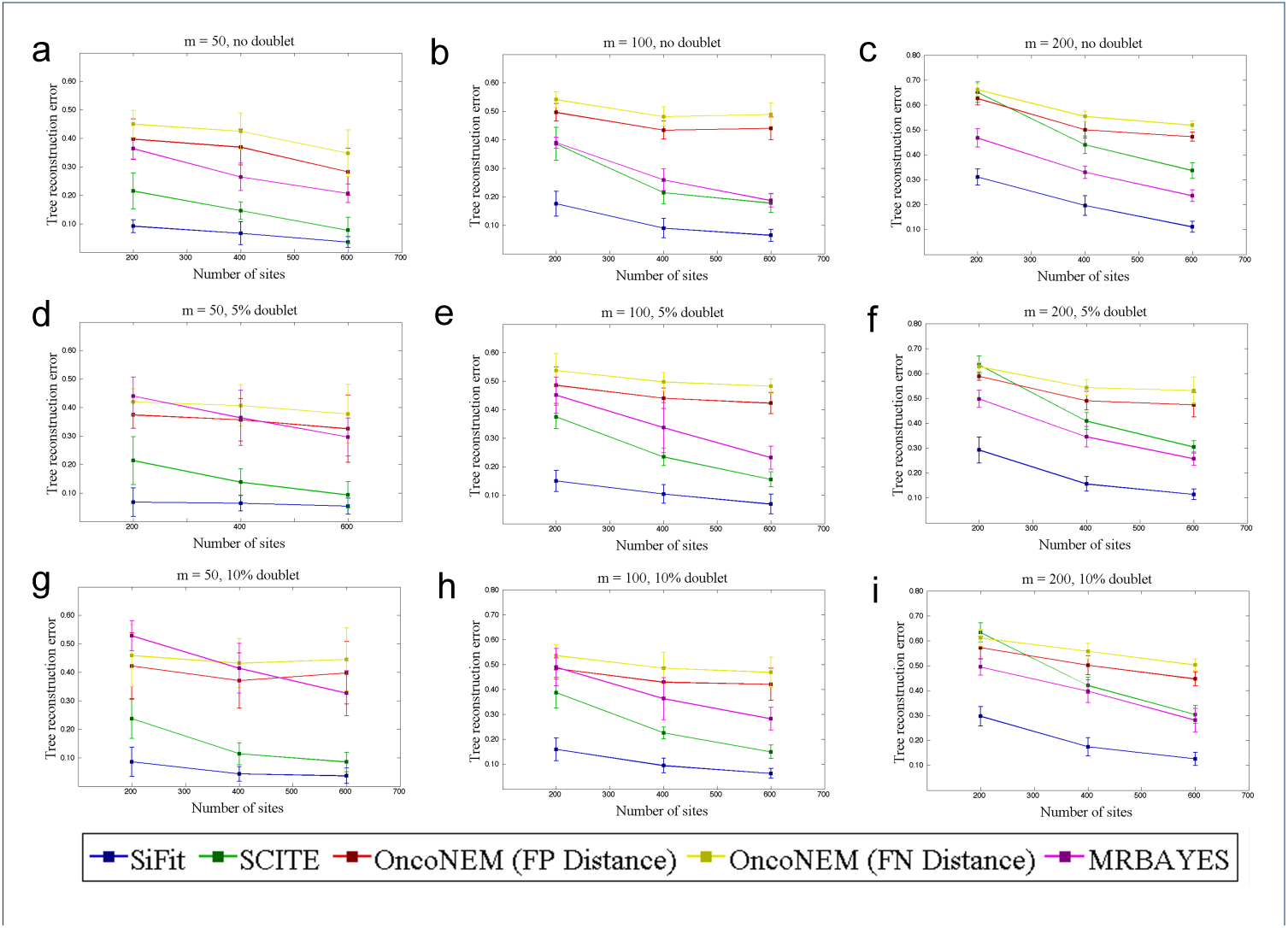
Performance comparison on datasets with varying number of cells. SiFit’s tree reconstruction accuracy is compared against that of SCITE, OncoNEM and MrBayes. The y-axis denotes the tree reconstruction error that measures the distance of the inferred tree from the ground truth. Three points for *n* = 200, *n* = 400 and *n* = 600 are plotted on the x-axis. In each case, the mean tree reconstruction error over 10 datasets is plotted. The vertical error bar indicates the standard deviation of tree reconstruction error over 10 datasets. **a** Performance comparison for datasets with 50 cells and *δ* = 0. **b** Performance comparison for datasets with 100 cells and *δ* = 0. **c** Performance comparison for datasets with 200 cells and *δ* = 0. **d** Performance comparison for datasets with 50 cells and *δ* = 0.05. **e** Performance comparison for datasets with 100 cells and *δ* = 0.05. **f** Performance comparison for datasets with 200 cells and *δ* = 0.05. **g** Performance comparison for datasets with 50 cells and *δ* = 0.1. **h** Performance comparison for datasets with 100 cells and *δ* = 0.1. **i** Performance comparison for datasets with 200 cells and *δ* = 0.1. For **d**-**i**, the tree reconstruction error is measured without considering the doublets in both true and inferred trees.

For the datasets with doublets, we measured tree reconstruction error in two ways: (1) doublets are removed from both the true tree and inferred tree and then the RF distance is calculated; and (2) RF distance is calculated between the true tree and inferred tree without any distinction of doublets. Since, doublets are hybrid of two cells that belong to two places in the tree, measuring the tree reconstruction error as in (1) ensures that position of all the other cells except the doublets are properly inferred, whereas, (2) measures the overall tree reconstruction error. Fig. 1 compares the algorithms in terms of tree reconstruction error as described in (1). SiFit outperforms the other three methods for all values of *δ*, *m* and *n*. SCITE and MrBayes’s performance are substantially affected by the presence of doublets, specifically for the datasets with smaller number of mutations. In comparison, SiFit’s performance is much robust in the presence of doublets while recovering the positions of the non-doublets in the tree. Even in terms of overall tree reconstruction error (measured as described in (2)), SiFit performs superior to other algorithms for all simulation settings corresponding to different values of *δ*, *n* and *m* (Additional file 1: Fig. S1).

### Inference with missing data

Due to uneven coverage and amplification bias, current single-cell sequencing datasets are challenged by missing data points where genotype states are unobserved. To investigate how missing data affect phylogeny reconstruction, we performed additional simulation experiments. For *m* = 100 and *n* = {200, 400, 600}, we generated datasets using the same error rates as before. For each combination of *δ*, *n* and *m*, we generated 10 datasets, for each of which, two other datasets with missing data = {10%, 25%} were generated. To generate the datasets with missing data, genotype information of sites were removed with probability 0.1, and 0.25 for missing data = {10%, 25%} respectively. SiFit’s results were compared against SCITE and OncoNEM, the results are shown in Fig. 2. For each value of *δ*, as the missing data rate increases from 0 to 25%, for each of the competing methods, we observe a steady increase in tree reconstruction error. For datasets without doublets (*δ* = 0), irrespective of the percentage of missing data, SiFit performs substantially better than SCITE and OncoNEM. SiFit’s likelihood calculation treats each missing data as contributing a marginal probability of 1, effectively making it equivalent to reducing the number of sites *n*. For the datasets with doublets, we measured tree reconstruction error in two ways as described in the previous section. SiFit outperforms both SCITE and OncoNEM irrespective of the way tree reconstruction error was measured (Fig. 2, Additional file 1: Fig. S2).

**Figure 2.**
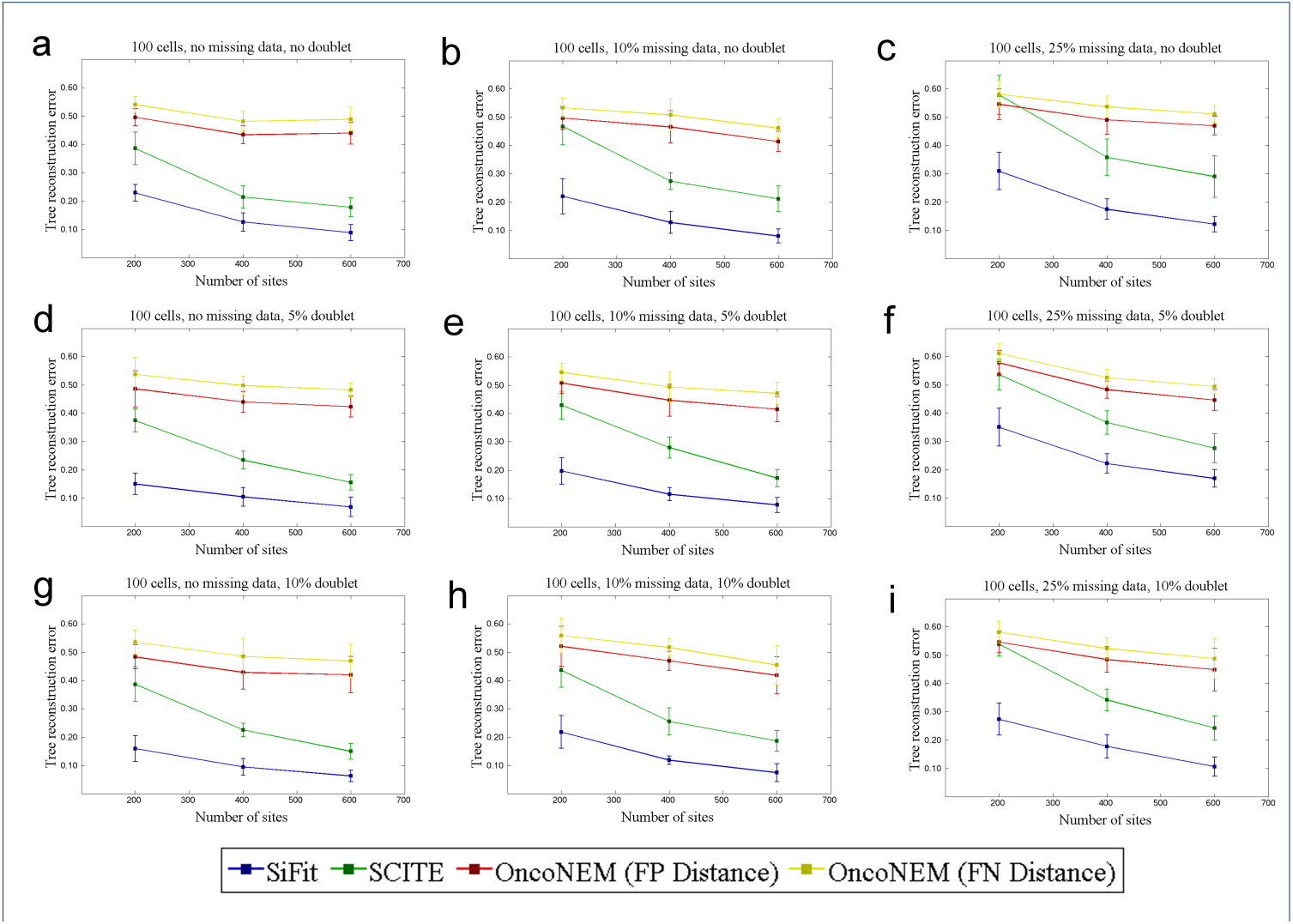
Performance comparison on datasets with missing data. SiFit’s tree reconstruction accuracy is compared against that of SCITE and OncoNEM on datasets with missing data. The y-axis denotes the tree reconstruction error that measures the distance of the inferred tree from the ground truth. Three points for *n* = 200, *n* = 400 and *n* = 600 are plotted on the x-axis. In each case, the mean tree reconstruction error over 10 datasets is plotted. The vertical error bar indicates the standard deviation of tree reconstruction error over 10 datasets. **a** Comparison for datasets without any missing data and *δ* = 0. **b** Comparison for datasets with 10% missing data *δ* = 0. **c** Comparison for datasets with 25% missing data *δ* = 0. **d** Comparison for datasets with without any missing data and *δ* = 0.05. **e** Comparison for datasets with 10% missing data *δ* = 0.05. **f** Comparison for datasets with 25% missing data *δ* = 0.05. **g** Comparison for datasets without any missing data and *δ* = 0.1. **h** Comparison for datasets with 10% missing data *δ* = 0.1. **i** Comparison for datasets with 25% missing data *δ* = 0.1. For **d**-**i**, the tree reconstruction error is measured without considering the doublets in both true and inferred trees.

### Robustness to increasing error rates

Allelic dropout is the major source of error in single-cell sequencing data resulting in false negatives [14]. To test the robustness of SiFit to increase in false negative rate, *β*, we simulated datasets with increased false negative rates. The number of cells, *m* was set to 100 and the number of sites, *n*, was set to 400. Mean false negative rate, *β*_*mean*_, was varied from 0.2 to 0.4 in steps of 0.1 i.e, *β*_*mean*_ ∈ {0.2, 0.3, 0.4}. The false negative rate of cell *c*, *β*_*c*_ was sampled from a normal distribution as described in the previous experiment. The false positive rate was set to *α* = 0.01. With these settings, for each value of *β*_*mean*_ *∈* {0.2, 0.3, 0.4}, 10 datasets were simulated for phylogeny reconstruction.

Performance of SiFit was compared against SCITE and OncoNEM. For different settings of false negative rates, SiFit consistently performs better than SCITE and OncoNEM by achieving the lowest tree reconstruction error (Fig. 3). For SCITE and SiFit, with the increase in the false negative rate, the tree inference error increases. For OncoNEM, the tree reconstruction error first increases and then decreases. The rate of increase in tree reconstruction error for SiFit is also much lower as compared to that of SCITE. This indicates SiFit’s higher robustness against amplification errors as compared to SCITE. OncoNEM’s tree reconstruction error is higher than SCITE and SiFit for all values of false negative rate. For OncoNEM, binarization of the data leads to loss of information and it employs a grid search to learn the parameters before learning the optimal tree. This divisive sequential approach of learning may lead to suboptimal solution if the initial solution gets stuck to local optima.

**Figure 3.**
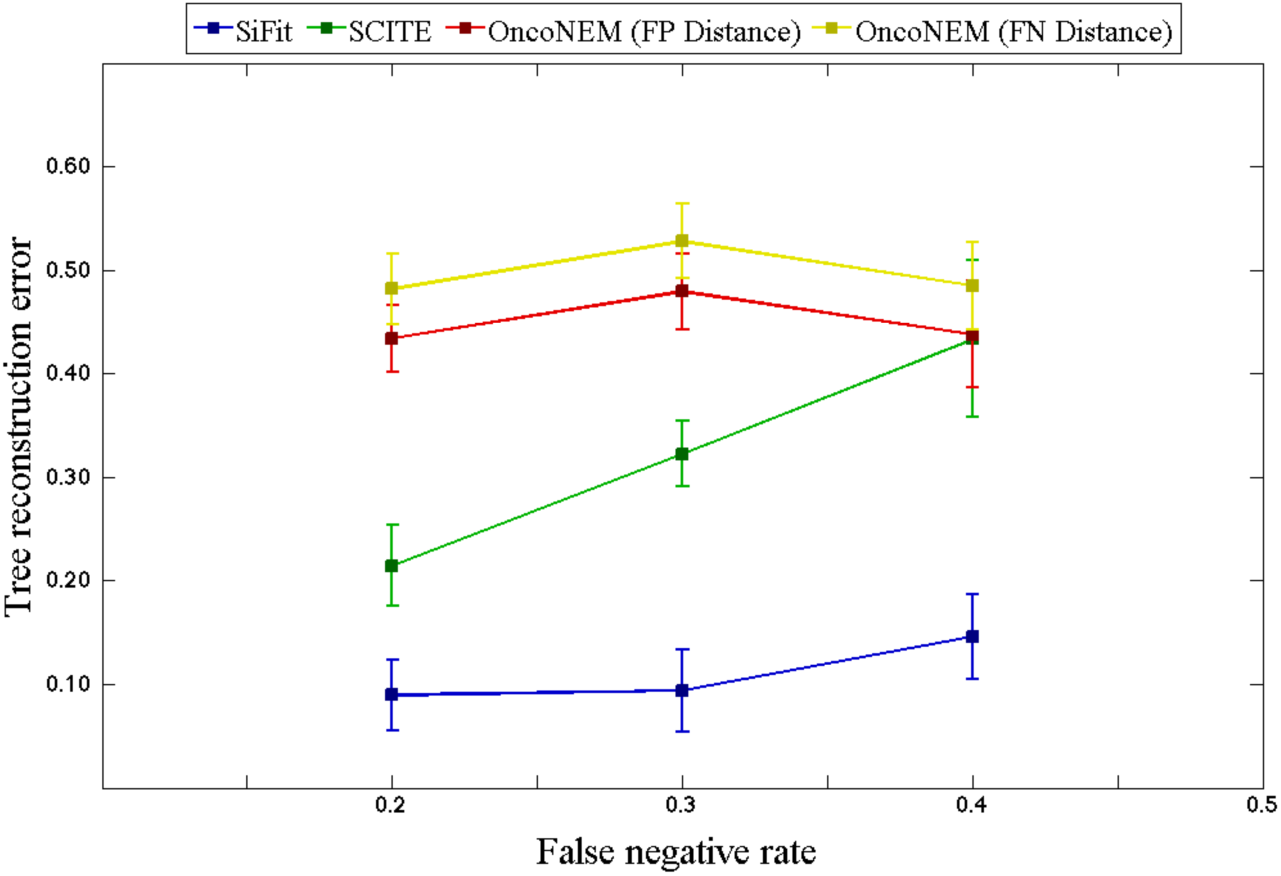
Effect of increase in error rates. SiFit’s tree reconstruction accuracy is compared against that of SCITE and OncoNEM for increasing false negative rate. The y-axis denotes the tree reconstruction error that measures the distance of the inferred tree from the ground truth. Four points corresponding to false negative rate, *β* = {0.2, 0.3, 0.4} are plotted. In each case, the mean tree reconstruction error over 10 datasets is plotted. The vertical error bar indicates the standard deviation of tree reconstruction error over 10 datasets.

### Estimation of error rates

In addition to the phylogenetic tree, SiFit also learns the error parameters from the data. To examine SiFit’s capability to estimate the false negative rate from data, we simulated 30 datasets from 30 random binary trees. For these datasets, the number of cells was set to 100 and the number of sites was set to 400 and the false positive rate was set to *α* = 0.01. The false negative rate, *β* was varied from 0.1 to 0.4. These imperfect data matrices were given to SiFit for inference of tree and false negative rate.

SiFit performed very well for estimating false negative rate as shown in Fig. 4. The maximum likelihood value of *β* learned from the data were highly correlated (0.9843) to the ones that generated the data. This experiment demonstrates SiFit’s ability to infer error parameters from data.

**Figure 4.**
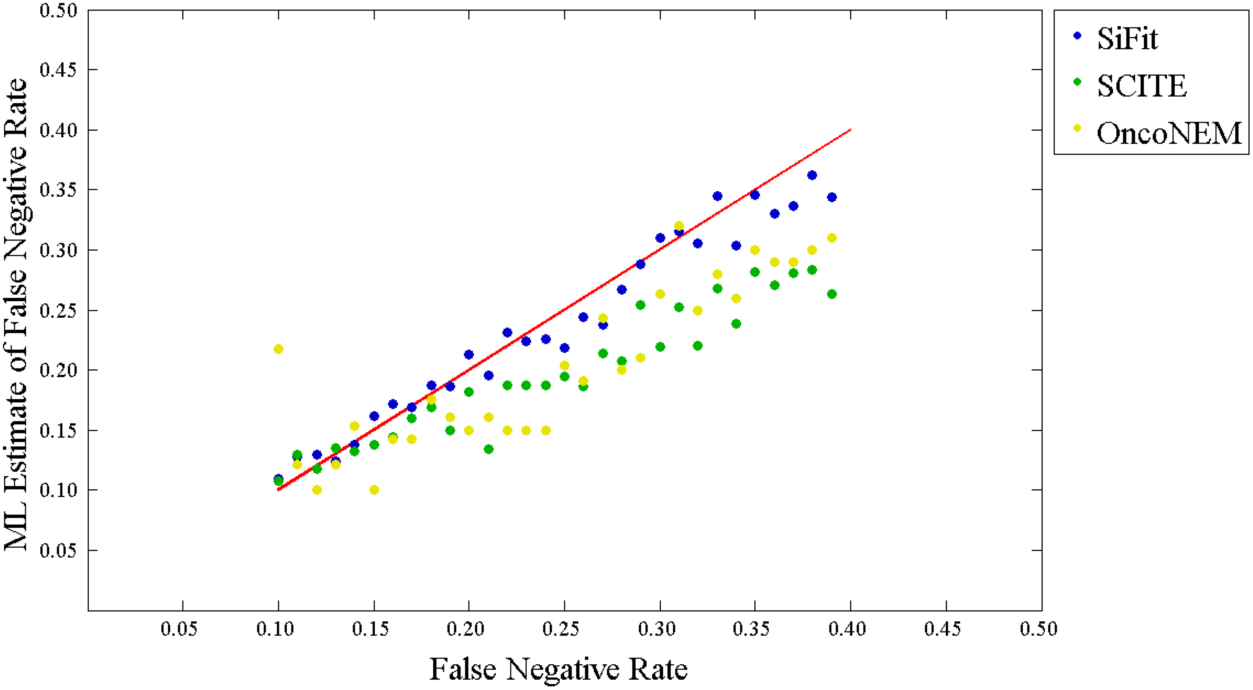
Estimation of error rates. The ML estimate of false negative rate is compared against the false negative rate used for generating the data. The red line represents the perfect estimate (correlation coefficient = 1). The blue dots represent the estimates by SiFit, the green dots represent the estimates by SCITE. The yellow dots correspond to estimates by OncoNEM.

SCITE and OncoNEM can also learn false negative rate from data. To compare SiFit’s estimate of error rate against that of OncoNEM and SCITE, we applied SCITE and OncoNEM to the same datasets for learning false negative rates. SCITE’s performance (correlation 0.9622) was better than that of OncoNEM (correlation 0.8766) but SiFit was the best performer. Specifically for datasets with higher false negative rate (*>* 0.2), SiFit’s estimates were much better than that of SCITE and OncoNEM. This indicates a degree of robustness of SiFit in the presence of higher error rates compared to the other methods.

### Runtimes

To measure the runtime of SiFit, we simulated datasets containing different number of cells. The number of cells, i.e, leaves in the trees, *m*, was varied as *m* = 100, *m* = 200 and *m* = 500. The number of sites, *n*, was varied as *n* = 200 and *n* = 400. The error rates were chosen as described in the previous experiments. For each combination of *m* and *n*, 10 datasets were simulated. For each of these datasets, SiFit was run for 200,000 iterations in a node with 24 CPU-cores (AMD 2.2 GHz). In each case, the average runtime for 200,000 iterations was recorded (Additional file 1: Fig. S3). For a fixed value of number of sites (*n*), with the increase in the number of cells in the tree, SiFit’s runtime increases almost linearly. This behavior is observed for both *n* = 200 and *n* = 400. This indicates that SiFit is scalable and will adopt well when the future experiments generate sequencing data consisting of thousands of single cells. The theoretical computational complexity of SiFit is described in the ‘Methods’ section.

#### Inference of tumor phylogeny from experimental SCS data

We applied SiFit to two experimental single-cell DNA sequencing datasets: exome sequencing from a non-herediatry colorectal cancer patient and high-throughput single-cell sequencing from a metastatic colorectal cancer patient. From these data we inferred the phylogenetic lineages of the tumor and ordered the chronology of mutations. These studies used different single-cell DNA sequencing methods and had different samples sizes and error rates, which we selected to show that SiFit is flexible and can be applied broadly to different single-cell mutation datasets.

### Phylogenetic lineage of adenomatous polyps and colorectal cancer

SiFit was applied to single-cell exome sequencing data from a non-hereditary colorectal cancer [46] patient. The dataset consisted of 61 single cells in total, with 35 cells were sampled from colorectal cancer tissue, 13 from an adenomatous polyp tissue and 13 from normal colorectal tissue. Variant calling resulted in the detection of 77 somatic SNVs from these 61 cells. In total, approximately 9.4% of the values were missing in the dataset. The reported genotypes were binary values, representing the presence or absence of a mutation at the SNV sites (Additional file 1: Fig. S4a).

To test whether the genotype matrix violates the “infinite sites assumption”, we ran the four-gamete test. The four-gametes theorem states that an *m × n* binary matrix, *M*, has an undirected perfect phylogeny if and only if no pair of columns contain all four binary pairs (0, 0; 0, 1; 1, 0 and 1, 1), where *m* represents the number of taxa (leaves of the tree) and *n* represents genomic sites [47]. The perfect phylogeny model conveys the biological feature that every genomic site mutates at most once in the phylogeny [47] and that mutations are never lost. The existence of perfect phylogeny shows that the data could fit the infinite sites model of evolution. A violation of four-gamete condition can indicate potential deviation from “infinite sites assumption”. However, it is important to note that for SCS data, There could be more than one potential events leading to violation of four-gamete test (see Additional file 1: Supplementary Note for more detail). The binary mutation matrix from this colorectal patient violated the four-gamete test, with 1847 (out of 2926) pairs of SNV sites that contained all four binary pairs.

The maximum likelihood tree inferred by SiFit on 77 SNVs is shown in Fig. 5. The tree shows that the normal cells are placed very close to the root. In the original study, some of the adenomatous polyp cells were found to have no somatic mutations and were speculated to have derived from normal colorectal cells. In the tree inferred by SiFit, these cells (ap8-ap13) are accurately placed along with the normal cells. The original study also reported a set of cells from the cancer tissue as normal cells because they did not contain any somatic mutations. The tree inferred from SiFit placed these cells along with the normal cells, representing a completely independent lineage that likely initiated from a different cell of origin. We performed k-medoids clustering using silhouette score (see “Method” for details) on the ML tree-based distance matrix. The cancer cells were clustered into two subpopulations (A and B). The chronological order of the mutations were inferred based on the inference of mutation status of the internal nodes. We extended (see “Method” for details) the algorithm in [48] for inferring ancestral sequences by accounting for single-cell specific errors. This enabled us to find the maximum likelihood solution for placing the mutations on the branches of the SiFit tree. 53 clonal mutations occurred in the trunk of the tree, including mutations in *LAMA1* (PI3K-Akt signaling pathway) and *ADCY3* (FGFR signaling pathway). These clonal mutations are driver events that likely led to the expansion of subpopulation A. The subpopulation B emerged from subpopulation A by acquiring additional subclonal mutations in *EPHA5, CASQ2 and SMARCE1*. The SiFit tree also shows the evolution of the adenomatous polyp cells (marked in blue), which evolved from the normal cells by acquiring mutations in *OR1B1* (GPCR signaling pathway), *DCDC5* and *MLLT1*. The adenomatous polyp cells evolved independently and further accumulated mutations in *CSMD1, FBXO15 and TCP11*. The tree inferred by SiFit represented the evolution of both the adenomatous polyp cells and the colorectal cancer cells and identified the order of the mutations that are associated with different signaling pathways and may have played a key role in the development of heterogeneity in this cancer patient.

**Figure 5.**
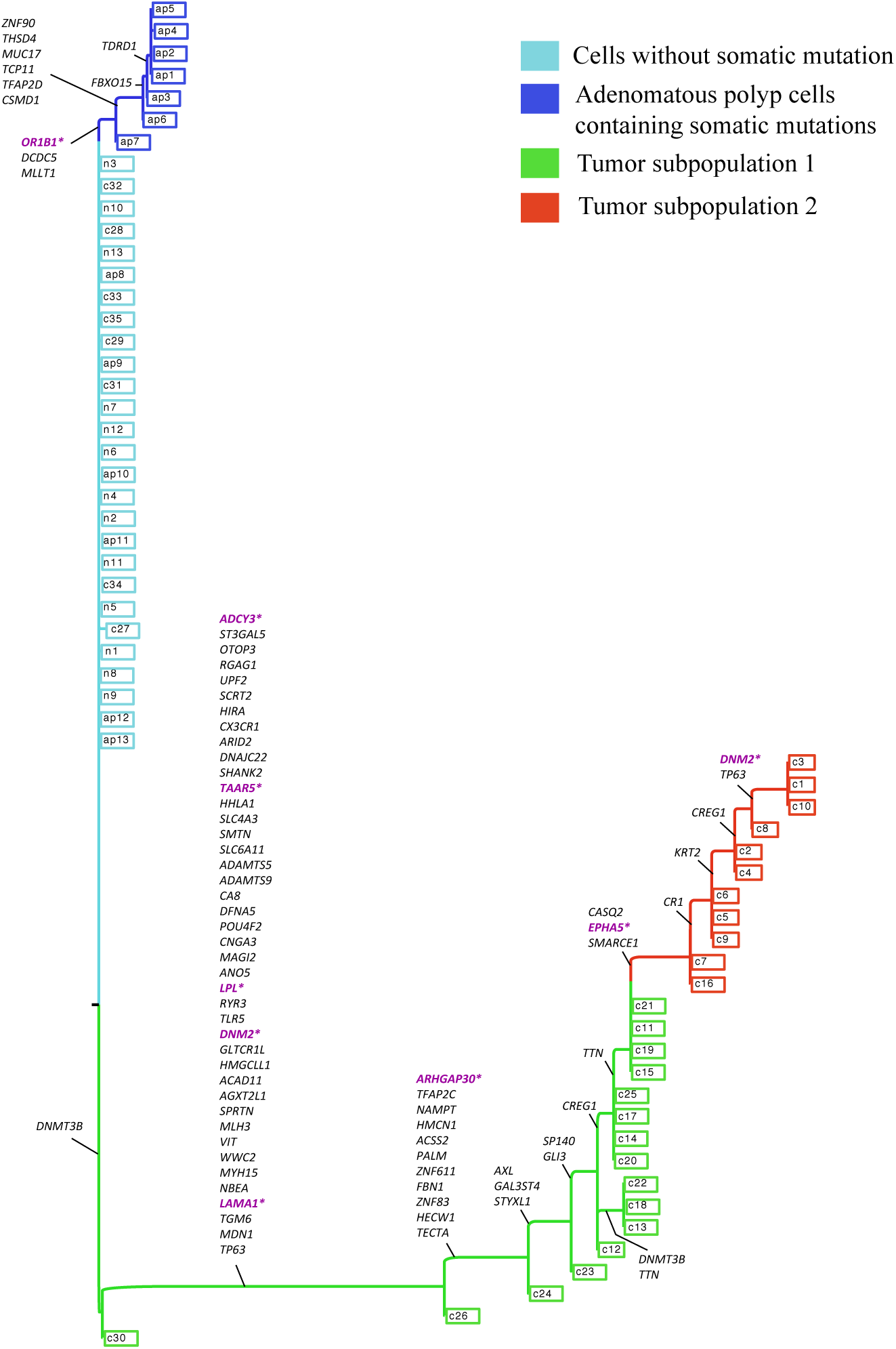
Maximum Likelihood phylogenetic tree reconstructed by SiFit for adenomatous polyps and colorectal cancer. The leaves with legend marked with ‘n’ are normal cells, leaves marked with ‘ap’ are adenomatous polyp cells, all other leaves are single tumor cells. Mutations are annotated on the branches of the tree. Important genes reported in the original study are marked in purple.

To compare the result of SiFit with other algorithms, we also applied SCITE and OncoNEM on this dataset. To enable direct comparison, SCITE was used to infer a binary leaf-labeled tree, which is a maximum likelihood solution with the single cells placed at the leaves of the tree. SCITE reported a single maximum likelihood (ML) tree (*T*_*SCIT E*_) from this dataset (Additional file 1: Fig. S5). We compared the tree inferred by SiFit (*T*_*SiFit*_) to the tree inferred by SCITE in terms of the likelihood value. Since, the ML tree inferred by SCITE (*T*_*SCITE*_) does not have branch lengths, we can not directly compute the likelihood value of *T*_*SCITE*_ using our likelihood function. Instead, we used the likelihood function of SCITE to compare the two trees. SCITE uses an expected mutation matrix defined by the mutation tree topology and sample attachments to compute the likelihood of a tree. After finding the maximum likelihood placement of the mutations on the SiFit tree (*T*_*SiFit*_), we obtained the expected mutation matrix, *E*, defined by *T*_*SiFit*_ and the annotated mutations on the branches of *T*_*SiFit*_ and then calculated its likelihood using Eq. 3 of [38]. This likelihood function of SCITE gives an edge to SCITE and is disadvantageous for SiFit because the branch lengths inferred by SiFit are ignored in this likelihood calculation. *T*_*SiFit*_ had a log-likelihood value of −632.5, which was substantially higher than the log-likelihood (-785.92) of *T*_*SCITE*_. This higher likelihood suggests that the tree inferred by SiFit explains the data better to that of SCITE on this experimental dataset.

We used OncoNEM to infer the cell lineage tree (*T*_*OncoNEM*_) from this dataset (Additional file 1: Fig. S6). OncoNEM can also estimate the occurrence of mutations on the cell lineage tree based on posterior probability. Since, OncoNEM follows “infinite sites assumption”, if a cell in the lineage tree contains a mutation, all its descendants should have that mutation. Based on this principle and OncoNEM’s estimate of the occurrence of mutations, we can compute an expected mutation matrix that is defined by the *T*_*OncoNEM*_. This enabled us to use the likelihood function of SCITE to compare *T*_*OncoNEM*_ against *T*_*SiFit*_. The log-likelihood value (-664.79) of *T*_*OncoNEM*_ was better than that of *T*_*SCITE*_ but it was worse than that of *T*_*SiFit*_. The higher likelihood of tree inferred by SiFit compared to that of OncoNEM and SCITE suggests that the expected mutation matrix defined by SiFit’s tree inferred under a finite-sites model of evolution explains the data better than the contemporaries inferred under “infinite sites assumption”.

### Phylogenetic lineage of a metastatic colorectal cancer patient

Next, we applied SiFit to infer the metastatic lineage of a colorectal cancer patient with a matched primary tumor and liver metastasis that was untreated. This dataset consisting of highly-multiplexed single-cell DNA sequencing data [31] from 178 single cells using a 1000 cancer gene panel. Variant calling resulted in the detection of 16 somatic SNVs from these 178 cells [49]. The false positive rate was estimated to be 1.52% and the false negative rate was estimated to be 7.89%. In total, approximately 6.9% of the values were missing in the dataset. The reported genotypes were binary values, representing the presence or absence of a mutation at the SNV sites (Additional file 1: Fig. S4b).

We ran the four-gamete test on this dataset, which identified 104 (out of 120) pairs of SNV sites violating the four-gamete test indicating potential violation of “infinite sites assumption”.

The maximum likelihood tree inferred by SiFit from this dataset is shown in Fig. 6. k-medoids clustering using silhouette score on the ML tree-based distance matrix identified three subpopulations of somatically mutated cells along with the population of cells without mutation. The subpopulation of cells (marked in cyan) without mutations consisted mostly of diploid cells, suggesting they are normal stromal cells. The first somatic subpopulation (marked in green) consisted of mostly diploid cells. The second subpopulation (matked in blue) consisted of mostly primary aneuploid cells and few diploid cells. The third subpopulation (marked in red) consisted of metastatic cells only. The chronological order of the mutations were inferred based on the maximum likelihood placement of the mutations on the branches of the tree. Three diploid cells in the first subpopulation first acquired a heterozygous nonsense mutation in *APC*. This mutation was present in all the descendants (all primary and metastatic tumor cells) suggesting that this was the first mutation that initiated the tumor. Subsequently, mutations were acquired in *KRAS* oncogene, *TP53* tumor suppressor gene and *CCNE1* oncogene respectively and lead to the expansion of the primary tumor mass. These primary tumor cells accumulated 7 additional somatic mutations. In later stages of the phylogeny, the accumulation of mutations in *EYS*, *ZNF521*, and *TRRAP* marked the point of metastatic divergence, after which tumor cells disseminated to the liver. Three more mutations occurred in *RBFOX1*, *GATA1* and *MYH9*. The phylogeny also indicates potential losses of mutations, including *POU2AF1*, which was lost in 17 primary tumor cells, and the mutation in *TCF7L2* that was lost in 4 metastatic tumor cells but these losses did not mark any point of divergence indicating they might be passenger mutations.

**Figure 6.**
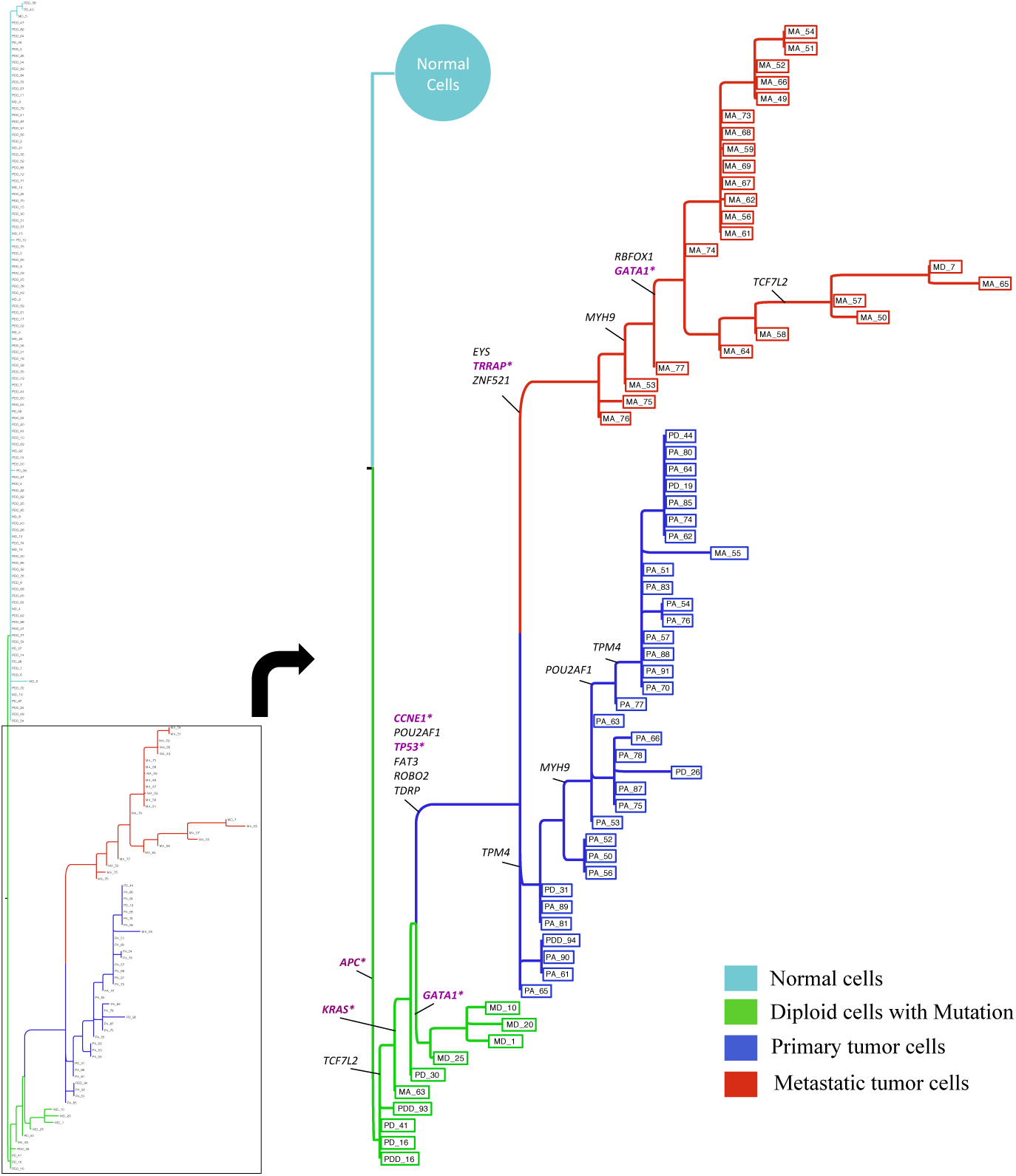
Maximum Likelihood phylogenetic tree reconstructed by SiFit for metastatic colorectal cancer patient. Mutations are annotated on the branches of the tree. The cancer genes and tumor-suppressor genes are marked in purple.

We also applied SCITE and OncoNEM on this dataset. SCITE inferred a single binary leaf-labeled tree (*T*_*SCITE*_, shown in Additional file 1: Fig. S7) which is the maximum likelihood solution with log-likelihood score of −387.68. To compute the likelihood of the tree (*T*_*SiFit*_) inferred by SiFit using SCITE’s likelihood function, we computed the expected mutation matrix *E* defined by *T*_*SiFit*_ using the maximum likelihood placement of the mutations on its branches. *T*_*SiFit*_ had a higher value of log-likelihood (−201.63) score. OncoNEM was used to infer a cell lineage tree (*T*_*OncoNEM*_, shown in Additional file 1: Fig. S8) from this dataset. We also estimated the occurrence of mutations on *T*_*OncoNEM*_ based on the posterior probability values. This enabled us to calculate the likelihood of *T*_*OncoNEM*_ through the computation of the expected mutation defined by *T*_*OncoNEM*_. *T*_*OncoNEM*_ had a log-likelihood value of −349.95, which is worse than *T*_*SiFit*_. The higher likelihood value of *T*_*SiFit*_ on this dataset suggests that the tree inferred by SiFit is superior to that of SCITE and OncoNEM in terms of explaining the data.

## Conclusions

Tumor phylogenies provide insight into the clonal substructure of tumors and the chronological order of mutations that arose during tumor progression. These lineages have direct applications in clinical oncology, for both diagnostic applications in measuring the amount of intra-tumor heterogeneity in tumors and for improving targeted therapy by helping oncologists identify mutations that are present in the majority of tumor cells. Single-cell DNA sequencing data provides an unprecedented opportunity to reconstruct tumor phylogenies at the highest possible resolution, however are challenged by extensive technical errors that are introduced during genome amplification. In this paper, we introduced SiFit, a probabilistic method for recreating the evolutionary histories of tumors under finite-site model of evolution from imperfect mutation profiles of single cells. This likelihood-based approach can infer the maximum likelihood phylogeny that best fits single-cell datasets with extensive technical noise. SiFit can also estimate the error rates of the single-cell DNA sequencing experiments. SiFit employs a resilient error model that can account for various technical artifacts in single-cell sequencing data, including allelic dropout (ADO), false positives and missing data. Our model is adaptable and can be easily extended to include position-specific error rates. SiFit also provides this flexibility in choosing the model of evolution, for which, we developed a finite-sites model of evolution that accounts for the effects of various events in tumor evolution such as point mutations, deletion, LOH, etc. in single-cell datasets. SiFit is robust to variation in error rates and performs consistently with varying number of cells in the dataset making it widely applicable to SCS datasets that vary in error rates and the number of cells sequenced.

The main difference of SiFit from existing methods, such as SCITE [38] and OncoNEM [37] is that SiFit introduces a finite-site model of evolution. Both SCITE and OncoNEM makes the “infinite sites assumption” that is frequently violated in cases of convergent evolution or reversal of genotypes, events that occur in human tumors due to LOH and chromosomal deletions [42]. SiFit also makes use of the high-resolution SCS data by utilizing the single cells as the taxonomic units of the reconstructed phylogenetic tree. On the other hand, SCITE reports a mutation tree, in which the lineage of the cells are not shown. OncoNEM reports a clonal tree, which is a condensed tree with multiple cells clustered into a clone. This type of clonal clustering and the use of clones as the taxonomic units, though useful for finding genealogical relationships between clones, is low-resolution as a clone represents a consensus of information from multiple single cells. The utilization of mutation information from each individual cell makes SiFit’s tree reconstruction method both robust and high-resolution.

SiFit performs accurately as evident from a comprehensive set of simulation studies that takes into account different aspects of modern SCS datasets by experimenting with varying number of cells in the dataset, wide range of error rates and different fractions of missing data. The simulation studies also demonstrated that SiFit substantially outperformed the state-of-the-art methods and is more robust to technical errors from WGA. We also applied SiFit to reconstruct the phylogeny for two experimental SCS tumor datasets from two patients with colorectal cancer, including one patient with a matched liver metastasis. SiFit accurately reconstructed the phylogenetic lineages of these tumors, and identified points in which subpopulations diverged from the main tumor lineages. These trees also provided insight into the order of mutations and the chronology in which they occurred during tumor progression.

SiFit’s phylogeny inference can potentially be improved by incorporating copy number variations along with single nucleotide variants. Recent studies [50] indicate that copy number follows a punctuated evolutionary model and are likely to provide insight into possible loss of heterozygosity (LOH) events and can facilitate in tree inference. Such an approach has previously been used in the context of bulk sequencing data [16] and can be incorporated for SCS data under a finite-site model of evolution. SiFit currently uses fixed error rates at every site. The error model can be further extended using position-specific error rates, where sites with lower-confidence mutations will have higher error rates and vice versa. The error model will have higher complexity in that situation and systematic model selection has to be performed. It is important to note that out of the three different types of events that could hint at deviation from “infinites sites assumption”, SiFit currently models events (deletions, LOH, etc.) that affect the same genomic site more than once and the FP and FN errors in SCS data. The other potential source, cell doublets, are not explicitly included in SiFit’s error model. To include doublets in the error model, it will be necessary to move beyond the phylogenetic tree to phylogenetic networks as the doublets are amalgamation of two separate genotypes and should be represented by a node of in-degree two. Another approach might be to treat them as nuisance parameter and integrate out during likelihood calculation.

As single-cell DNA sequencing becomes more high-throughput [31, 51] enabling hundreds of cells to be analyzed in parallel at reduced cost and throughput, SiFit is poised to analyze the resulting large-scale datasets to understand the evolution of clones during tumor progression. SiFit represents a major step forward in understanding the tumor phylogeny from SCS data and will have important translational applications for improving cancer diagnosis, treatment and personalized therapy [14, 52]. Although the current study is focused on cancer, SiFit can potentially also be applied to single-cell mutation profiles from a wide variety of fields including immunology, neurobiology, microbiology and tissue mosaicism [53]. These applications are expected to provide new insights into our understanding of cancer and other human diseases.

## Methods

### Input data

The input to SiFit is a matrix *D*_*n×m*_ = (*D*_*ij*_) of observed genotypes, where *i* ∈ {1, …, *n*} denotes the index of genomic locus, *j* ∈ {1, …, *m*} is the index of the single cell and *D*_*ij*_ is the observed genotype at the *i*^*th*^ site of cell *j*. The genotype matrix can be binary or ternary depending on the representation of the data. For binary matrix, *D*_*ij*_ ∈ {0, 1, *X*}, where 0, 1 and *X* denote the absence of mutation, presence of mutation and missing data respectively. For a ternary matrix, *D*_*ij*_ can take value from the set, {0, 1, 2*, X*}, where 0 denotes homozygous reference genotype, 1 and 2 denote heterozygous and homozygous non-reference genotypes respectively and *X* denotes missing data.

### Model of single-cell errors

False positive errors and false negative errors are the two different types of noises that could be present in the genotype matrix. If *α* is the false positive error rate and *β* is the false negative error rate, then for a ternary genotype matrix, the relationship between the true and observed genotype matrices is given by:

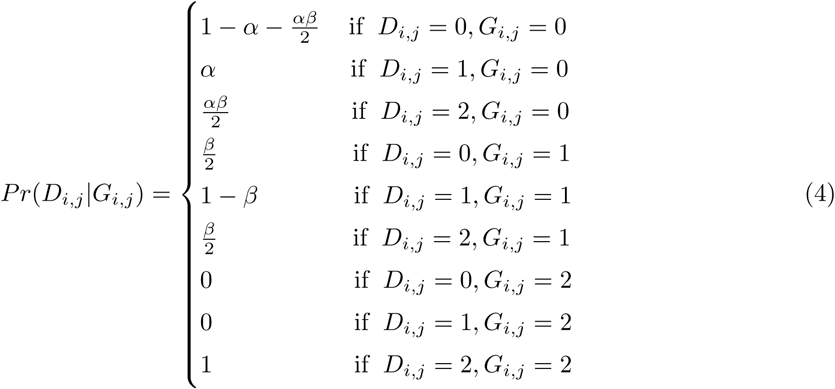

where *G*_*i,j*_ is the unobserved true genotype at the *i*^*th*^ site of cell *j*. A true homozygous non-reference genotype (site with true homozygous mutation) is affected by neither false positive error nor allelic dropout. A false negative error can affect the heterozygous genotype and combined with false positive error can also affect homozygous reference genotype. False positive error can affect homozygous reference genotypes.

Single-cell datasets also contain missing data, sites for which genotype information is missing. In our computation, we take *P r*(*D*_*i,j*_*|G*_*i,j*_) = 1 whenever *D*_*i,j*_ = *X*. By doing so, we marginalize the effect of missing data over three possible true genotypes and is reflected in the likelihood computation.

### Likelihood of a phylogenetic tree

#### Phylogenetic tree

We consider that the phylogenetic tree for single cells is a rooted directed binary tree *𝒥* = (*T*, **t**). It has two components, a tree topology *T* and a vector of branch lengths **t**. The phylogenetic tree represents the genealogical relationship among a set of single cells. The root of this tree has homozygous reference genotypes at all sites. The leaves of the tree represent the observed single cells. The internal nodes represent ancestral cells that are not observed in the data. Cells evolve along the branches of the tree following a model of evolution and the branch length denotes expected number of mutations per site.

#### Model of evolution

The finite-site model of evolution for SCS data (*ℳ*) is modeled using a continuous-time Markov chain that assigns a probability with each possible transition of genotypes. We assume that the genomic sites evolve identically and independently. Assuming three possible genotype states {0, 1, 2} (for ternary data) for a genomic site, the model of evolution can be represented by a 3 × 3 transition probability matrix. The transition probability matrix, *P*_*t*_, along a branch of length *t* is computed by matrix exponentiation of the product of the transition rate matrix (*Q*) of the Markov chain and the branch length. The entries in the transition rate matrix denotes the infinitesimal rates (during infinitesimally small time, Δ*t*) at which the continuous-time Markov chain moves between genotype states. We also consider that the time Δ*t* is the smallest unit of time during which only one event can occur at a site. Since, we are considering the somatic mutation sites, the infinitesimal rate for the genotype transition 0 → 1 is set to 1. This accounts for the point mutations. LOH events can result in the genotype transitions 1 → 0 and 1 → 2 whereas deletions can result in the genotype transitions 1 → 0, 1 → 2 or 2 → 1. To compute the infinitesimal rates for these transitions, we introduce two parameters *λ*_*d*_ and *λ*_*l*_ that account for the effects of deletion and LOH respectively. The product of the transition rate matrix and the branch length (*t*) is given by:

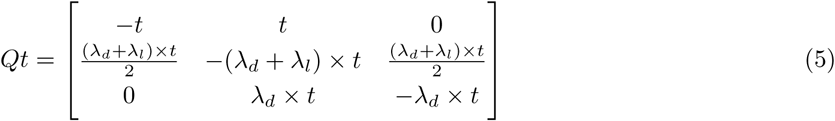

In Eq. (5), *Qt*(*i, j*) denotes the rate of genotype *i* changing to genotype *j* along a branch of length *t*, *i, j* ∈ {0, 1, 2}. *λ*_*d*_ and *λ*_*l*_ constitute the set of parameters (*ℳ*_*λ*_) of the model of evolution.

The transition probability matrix, *P*_*t*_ is given by:

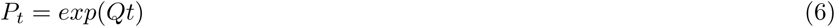

*P*_*t*_(*i, j*) denotes the probabality of transition of genotype *i* to genotype *j* along a branch of length *t*. Each entry of *P*_*t*_ is a function of *t*, *λ*_*d*_ and *λ*_*l*_.

For binary genotype states, the product of transition rate matrix and branch length is given by:

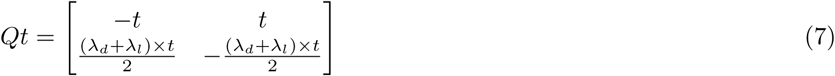

and the transition probability matrix is computed using Eq. (6).

#### Likelihood

Since we assume that each site evolves independently and the technical errors affect each site independently, the likelihood for the observed genotype matrix given a phylogenetic tree *𝒥*, error rates, ***θ*** and parameters of model of evolution, *ℳ*_*λ*_ is given by

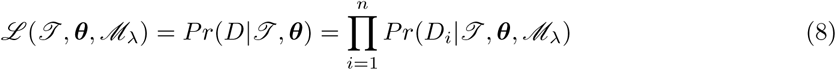

where *D*_*i*_ is the observed data at site *i* and it is a vector with *m* values corresponding to *m* single cells. Let *γ* be the set of possible genotypes. If *v* be an internal node of the tree with children *u, w*, and let 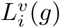 denote the partial conditional likelihood defined by

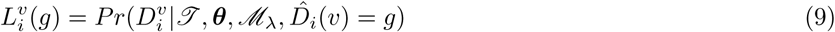

where 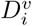 is the restriction of data *D*_*i*_ to the descendants of node *v* and 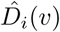 is the ancestral genotype for *i*^*th*^ site at node *v*. 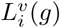 is the likelihood at site *i* for the subtree rooted at node *v*, given that the genotype at *v* is *g*.

The likelihood of the complete observed data *D*_*i*_ at the *i*^*th*^ site is given by:

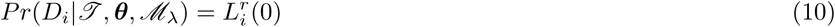

where *r* is the root of the tree and since we consider that the genotypes at the root are all homozygous reference (0), the probability 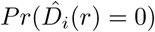 equals 1. The partial conditional likelihood function satisfies the recursive relation

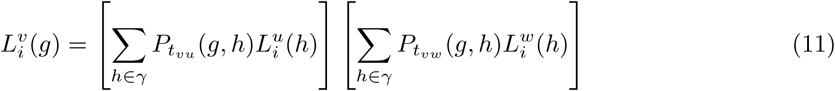

for all internal nodes *v* with children *u* and *w*. *t*_*vu*_ and *t*_*vw*_ are the branch lengths corresponding to branches that connect *v* to *u* and *w* respectively. 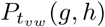 and 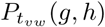 are the transition probabilities that are calculated using Eq. (6) with argument *t*_*vu*_ and *t*_*vw*_ respectively. For a leaf of the tree that denotes single cell *j*, the partial likelihood is given by

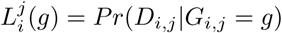

where *P r*(*D*_*i,j*_*|G*_*i,j*_) is calculated using either Eq. (2) or Eq. (4) depending on the data. The partial likelihood values at the leaves are computed based on the error rates of SCS data.

The log-likelihood for the observed genotype matrix given a phylogenetic tree *𝒥* error rates, ***θ*** and model parameters, *ℳ*_*λ*_ becomes a summation over *n* sites as in Eq. (12)

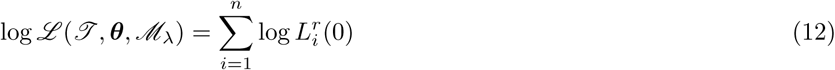

This likelihood computation uses Felsenstein’s pruning algorithm [54] for calculating the likelihood of a phylogenetic tree with the transition probabilities given by Eq. (6). For the calculation of the partial likelihoods for leaves, we use the SCS error model instead of values suggested in [54].

### Search algorithm to infer phylogeny

We developed a heuristic search algorithm to stochastically explore the joint space of phylogenetic trees, error rates and evolution model parameters. In the joint (*𝒥*, ***θ****, ℳ*_*λ*_) space, we need to consider three different types of moves to propose a new configuration. In tree changing moves, a new phylogenetic tree, *𝒥’* is proposed from current state *ℳ*. In error rate changing moves, a new error rate, ***θ*** *’* is proposed from current error rate ***θ***. In parameter changing modes, a new value of the parameter, *ℳ*_*λ’*_ is proposed from the current parameter value *ℳ*_*λ*_. If the proposed configuration results in a higher likelihood, it is accepted, otherwise rejected.

With a small probability, the proposed configuration is accepted or rejected based on an acceptance ratio (only for tree changing or error rate changing moves). The acceptance ratio for proposing a new phylogenetic tree is given by,

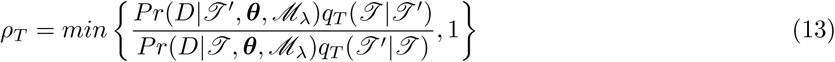

which involves calculating the ratio of the likelihood of the new configuration and the current configuration. Acceptance ratio also requires a proposal ratio which is computed based on *q*_*T*_, the proposal distribution for proposing new tree. A new error rate ***θ*** *’* is accepted with ratio given by,

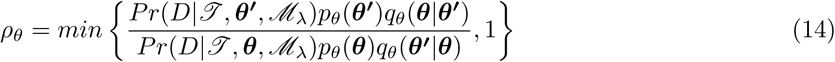

which takes into account the ratio of likelihoods of new and current configurations, the ratio of prior probability of new and current error rates and also a proposal ratio. *p*_*θ*_ is the prior distribution on error rate and *q*_*θ*_ is the proposal distribution for proposing new error rate. These steps of the search heuristic are motivated by Metropolis-Hastings algorithm [55] for doing Markov Chain Monte Carlo (MCMC) sampling and helps in better exploration of the likelihood space. The inference algorithm is shown in Algorithm 1

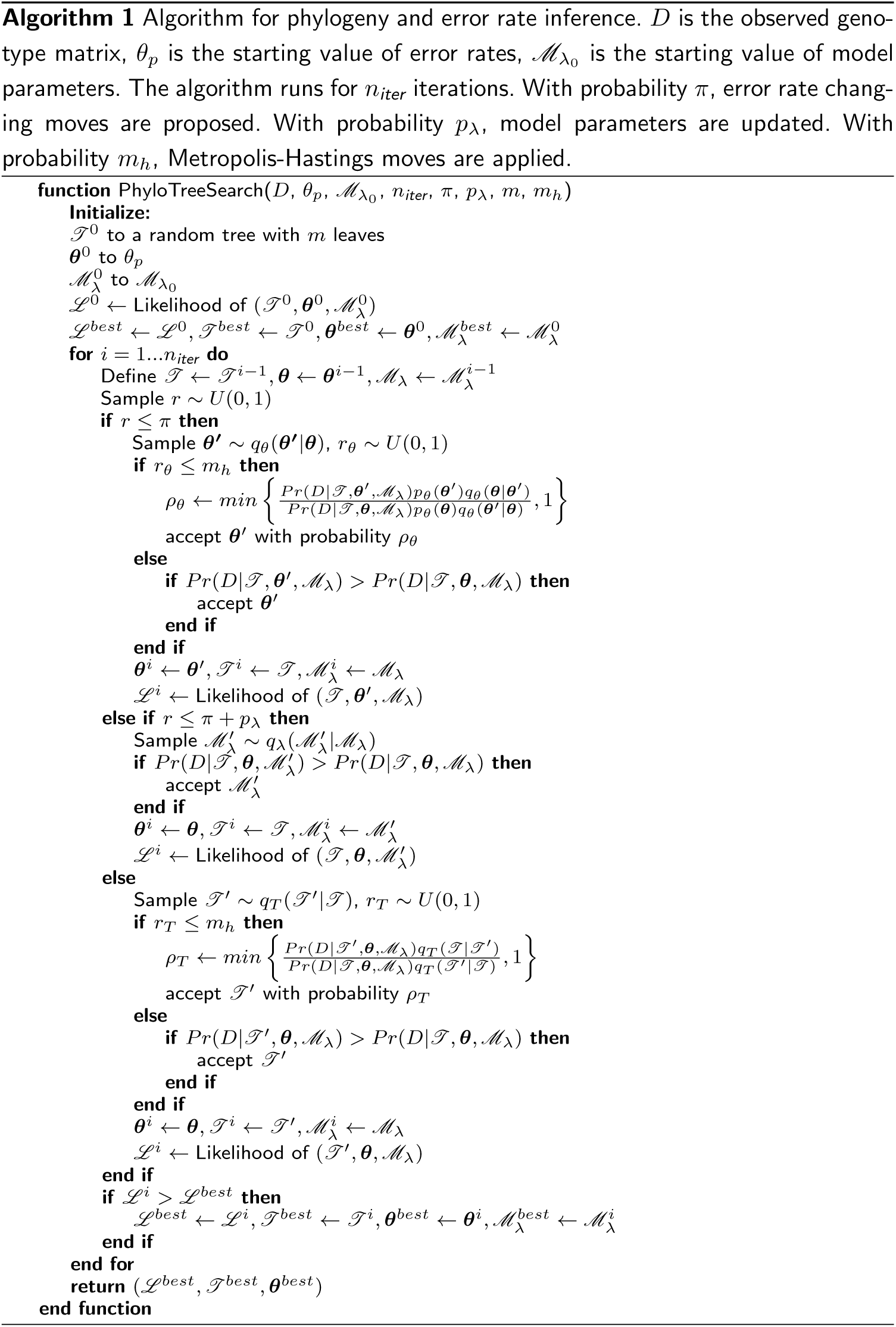

### Tree proposals

To explore the space of trees we need efficient moves that can make small and big changes in the tree topology. Also, we need moves that change only the branch lengths instead of changing the topology. To ensure that our search does not get stuck to a local optimum, we use a combination of different types of moves. Lakner et al. [56] described several tree proposal mechanisms that are effective in Bayesian phylogenetic inference. Since our goal is to effectively search the tree space, we can employ the same tree proposals in our search algorithm. We adopt two different types of tree proposals described in [56] in our search process, branch change proposals that alter branch lengths and branch-rearrangement proposals that alter the tree topology. The branch-rearrangement proposals can be divided into two subtypes: the prune and reattach moves and the swapping moves.

For proposing new branch length, we draw a sample *u* from a uniform distribution on [0, 1) and then get a random number *r*^*\**^ by applying the transformation *r*^*\**^ = e^*η*(*u-*0.5)^. The new branch length *l*^*\**^ is a product of current branch length *l* and *r*^*\**^. In this way, we update branch length of all branches. This ensures that branch lengths are locally changed the proposal ratio becomes a product 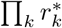, where *k* is the total number of branches in the tree. *η* is a tuning parameter that is set to the value suggested in [56].

We consider two types of pruning-regrafting moves, namely Random Subtree Pruning and Regrafting (rSPR) and Extending Subtree Pruning and Regrafting (eSPR), which were described in [56]. The pruning-regrafting moves randomly select an interior branch, prune a subtree attached to that branch, and then reattach the subtree to another regrafting branch present in the other subtree. For rSPR, the regrafting branch is chosen randomly. For eSPR, an extension probability guides the movement of the point of regrafting across one branch at a time. The eSPR move favors local rearrangements.

We consider three types of swapping moves, namely Stochastic Nearest Neighbor Interchange (stNNI), Random Subtree Swapping (rSTS) and Extending Subtree Swapping (eSTS). stNNI chooses an internal branch as the focal branch and stochastically swaps the subtrees attached to the focal branch. eSTS also involves the swap of two subtrees but not necessarily nearest neighbors. The subtrees are chosen according to an extension mechanism similar to eSPR. For rSTS, two randomly chosen subtrees are swapped.

At each step of the search algorithm, one of these six moves is chosen with a fixed probability. The proposal ratio associated with each branch-rearrangement proposal is described in detail in [56].

### Estimation of error rate

During the search process, we also update error rates. The estimates of error rates that are input to SiFit are used to design the prior probability *p*(*θ*). The error rate being a probability (value between 0 and 1), we choose a beta prior. The mean of the prior is estimated from the input error rate and observed genotype matrix. We choose a large standard deviation to cover a wide range of values. We choose a normal distribution as the proposal distribution for proposing new error rate. At each generation, the normal distribution is centered on the current value of error rate. A user specified fixed probability determines whether, in a particular iteration, a new error rate will be proposed.

### Estimation of parameters of model of evolution

The parameters of the model of evolution, *λ*_*d*_ and *λ*_*l*_ are also updated during the search process. For each of these parameters, the next value is proposed from a normal distribution centered at the current value. The standard deviation is chosen so that a wide range of values are covered. These parameters being relative quantities (denote rates of deletion and LOH respectively relative to the rate of point mutations), we choose a beta distribution as their prior. Similar to proposing new error rates, a user specified fixed probability determines whether, in a particular iteration, a new value of these parameters will be proposed.

### Complexity analysis

In each step of the algorithm, finding the likelihood of the tree is the most expensive task. For *m* single cells, *n* sites, the likelihood calculation takes *𝒪*(*mk*^2^*n*), where *k* is the maximum number of states per site. For genotype data, *k* = 3 and for binary mutation matrix, *k* = 2. The number of iterations used in SiFit is user-defined. Assuming *i* to be the number of iterations used for running SiFit, the overall complexity becomes *𝒪*(*mk*^2^*ni*).

#### Tree inference error metric

To measure the accuracy of tree inference, we used a metric that compares the topology of the inferred tree to that of the true tree and computes a distance between the two. This metric on general phylogenetic trees was proposed in [45] and it is based on the symmetric difference between the bipartitions of the two trees. The topology of a tree can be represented by the bipartitions present in the tree. A bipartition of a tree based on an edge gives us two set of leaves that would be formed by deleting the edge. If *ℰ* is the set of edges of *𝒥*, then the bipartition encoding of *𝒥*, denoted by *C*(*𝒥*) = {*ξ*(*e*) : *e ∈ ℰ*}, is the set of bipartitions defined by each edge in *𝒥*. *ξ*(*e*) is the bipartition on the leaf set of *𝒥* produced by removing the edge *e* from *𝒥*. We consider three distances between two trees.

If *𝒥*_*t*_ is the true tree on a set of single cells *𝒮* and *𝒥* _*i*_ is the inferred tree, then the following are the three inference error metrics.

*False Negative (FN) distance*, This counts the edges in *𝒥*_*t*_ that induce bipartitions that are not present in *C*(*𝒥*_*i*_). This distance is normalized by dividing by the total number of bipartitions in *𝒥* _*t*_, i.e. 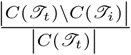

*False Positive (FP) distance*, This counts the edges in *𝒥*_*i*_ that induce bipartitions that are not present in *C*(*𝒥*_*t*_). This distance is normalized by dividing by the total number of bipartitions in *𝒥*_*i*_, i.e. 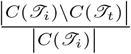

*Robinson-Foulds (RF) distance*. The Robinson-Foulds distance is the average of FP and FN distance. This is the most common error metric.

If the two trees to compare are binary then we use RF distance between them as the error metric. For binary trees, FP, FN and RF distances are equal to each other. To compare a true binary tree to an inferred non-binary tree, we compute FP and FN distances separately.

SiFit, SCITE and MrBayes output binary tree which can be compared against the true tree in terms of RF distance. For OncoNEM, we consider the cell lineage tree that it infers and then we convert the cell lineage tree to an equivalent phylogenetic tree by projecting the observed single cells to leaves (shown in Additional file 1: Fig. S8). The equivalent phylogenetic tree might be binary or non-binary and we compute both FP and FN distances for it when comparing to the true tree.

#### Inference of ancestral sequences and order of mutations

The inference of the chronological order of mutations in the tumor lineage requires the inference of mutation status of the internal nodes so that the mutations can be placed on the branches of the phylogeny. We infer the mutational profiles of the internal nodes using a likelihood-based approach that finds the most likely mutational profile for an internal node given the phylogenetic tree and error rates. We extend the dynamic programming algorithm for inferring ancestral sequences described in Pupko *et. al* [48] to account for the error rates of the single cells.

For a single cell *c* at the leaf of the tree, the partial likelihood for a genotype *g* at site *i* is calculated as 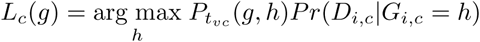 and mutation state, *m*_*c*_(*g*), is set to *h* that attains the maximum value for partial likelihood. *v* is the parent of *c* and *t*_*vc*_ is the branch length connecting *v* to *c*. For a missing data, *P r*(*D*_*i,c*_*|G*_*i,c*_ = *h*) becomes 1. For a nonroot internal node, *u*, with children *y* and *z*, the partial likelihood is calculated as 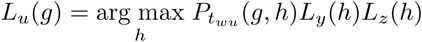 and the mutation state, *m*_*u*_(*g*), is set to *h* that attains the maximum value. For the root of the tree, mutation state *m*_*r*_ = 0 and the mutation state for an internal node, *u*, whose parent *w*’s mutation state is already determined as *g*, is chosen as *m*_*u*_(*g*).

After inferring the mutational profiles of the internal nodes, the mutations on a branch can be found by finding the SNV sites for which the mutational status of the two nodes at the two ends of the branch differ.

#### Clustering of cells

To cluster the cells into subpopulations for the tumor datasets, we used k-medoids clustering with silhouette scores. A distance matrix was obtained for the cells containing mutations from the ML tree reconstructed by SiFit, in which, an entry represents the distance between two cells. The distance between two cells was calculated by summing the branch lengths on the path that connects the two cells. K-medoids clustering was performed on the resulting distance matrix using ‘clustering’ library of R (http://www.r-project.org) and the number of clusters was varied from 2 to 5. In each case, the average silhouette score was measured and the number of clusters that maximized silhouette score was reported as the optimal number of clusters.

#### Simulation of synthetic data

### Evolution of single-cell sequences

To simulate single-cell datasets, first, a random binary tree is constructed on a leaf set of single cells by a recursive algorithm that randomly divides the set of cells into two subtrees that are also randomly generated, and then joins them into a single tree by choosing a root that has the two subtrees as the left and right child.

**Table 1.**
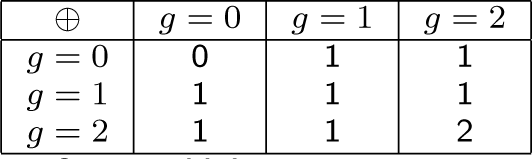
Expected genotype state after combining two genotypes using the binary operator ⊕.

We specify the number of sites, *n* in the single-cell genome. The root node of the phylogeny is populated with homozygous reference genotype (*g* = 0) at each site. In each branch of the tree, a Poisson distributed number of sites, *p*, are mutated. If *t* is the branch length, the parameter for the Poisson distribution is chosen as *t × n*, so that on an average, a child node in the tree differs from its parent by a proportion of loci which is given by the branch length. When mutating a new site, the genotype changes from homozygous reference (*g* = 0) to heterozygous (*g* = 1). Recurrent mutations are introduced with probability *r*. If the locus in the node, for which a recurrent mutation happens, has a homozygous reference genotype (*g* = 0), then a parallel mutation happens in that branch, i.e, the genotype changes from homozygous reference (*g* = 0) to heterozygous (*g* = 1). If the locus in the node already contains a mutated genotype then a back mutation results in reverting the genotype to homozygous reference (*g* = 0). To simulate loss of heterozygosity (LOH) events, the loci with heterozygous (*g* = 1) genotypes are set to either homozygous reference (*g* = 0) or homozygous non-reference (*g* = 2) genotypes with probability *ω*. If LOH happens at a locus, either of the homozygous genotypes are chosen with equal probability. Deletion is simulated with probability *d* at a branch. Deletion can affect multiple loci at a time. For a heterozygous site, deletion can happen for any of the copies resulting in either of the homozygous genotypes (*g* = 0 or *g* = 2). Deletion does not affect the homozygous reference genotypes but can change the homozygous non-reference genotypes to heterozygous genotype. In this way, sites are evolved at each branch of the tree. At the corner case, when there is no new locus to mutate at a branch, recurrent mutations are introduced. After considering all the branches of the tree, we have the single cell genotypes at the leaves of the tree.

### Simulating doublets

Doublets are events when two cells get trapped in the same well resulting in merging the genotypes of the two cells. To model doublets we need to define the expected genotype state which is a combination of two genotype states. The expected genotype state can be defined by a binary operator *⊕* whose results for SNV data are shown in Table 1. *δ* denotes the fraction of cells that are doublets. With probability *δ*, a cell is chosen to be a doublet and its genotype is combined with that of a randomly sampled co-trapped cell (genotype of which is the copy of another cell in the tree) to form the new genotype as defined by the *⊕* operator. The pseudocode for simulating doublets is shown in Algorithm 2.

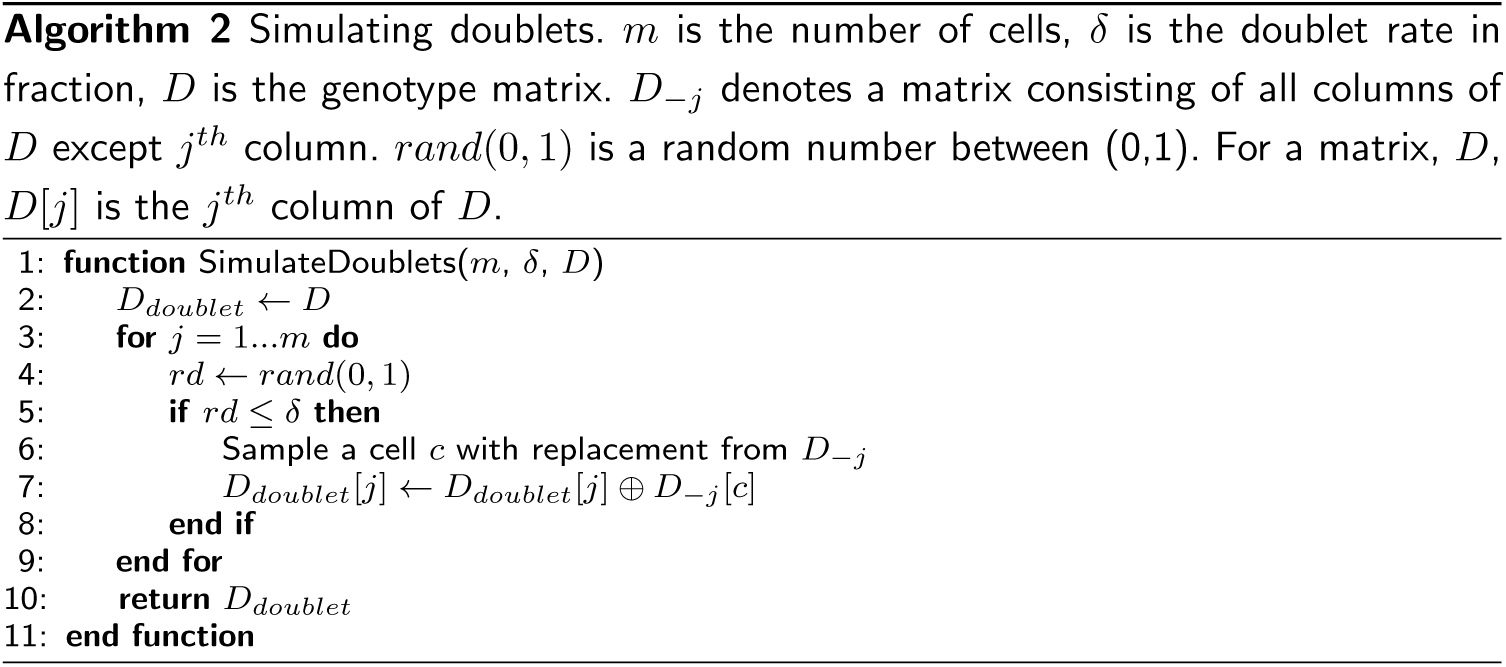

#### Software availability

SiFit has been implemented in Java and is freely available at https://bitbucket.org/hamimzafar/sifit. This implementation uses PhyloNet [57] and Habanero-Java library [58].

## Availability of data and materials

Raw sequencing data for the human tumor datasets that are analyzed have been deposited in the SRA database, under SRA numbers SRP067815 and SRP074289 respectively. The genotype matrix for the non-hereditary colon cancer patient has been reproduced from Fig. 3a of [46]. The genotype matrix for the metastatic colon cancer patient has been reproduced from Supplementary figure 4a of [49]. Both the data matrices are available with SiFit.

### Ethics approval

Not applicable.

### Author’s contributions

HZ, NN, KC and LN designed the study. HZ and AT developed the algorithm and implemented the software. HZ ran the experiments. All authors wrote and approved the manuscript.

### Funding

The study was supported by the National Cancer Institute R01 CA172652 (K.C.), cancer center support grant P30 CA016672 and Andrew Sabin Family Foundation.

## Acknowledgements

The authors thank Sri Raj Paul for providing useful suggestion regarding parallelizing part of the source code.

## Additional Files

Additional file 1 — Supplementary Material

This file contains supplementary note and supplementary figures.

## References

1. Nowell, P.: The clonal evolution of tumor cell populations. Science 194(4260), 23–28 (1976)

2. Merlo, L.M.F., Pepper, J.W., Reid, B.J., Maley, C.C.: Cancer as an evolutionary and ecological process. Nat Rev Cancer 6(12), 924–935 (2006)

3. Pepper, J.W., Scott Findlay, C., Kassen, R., Spencer, S.L., Maley, C.C.: Synthesis: Cancer research meets evolutionary biology. Evolutionary Applications 2(1), 62–70 (2009)

4. Yates, L.R., Campbell, P.J.: Evolution of the cancer genome. Nat Rev Genet 13(11), 795–806 (2012)

5. Greaves, M., Maley, C.C.: Clonal evolution in cancer. Nature 481(7381), 306–313 (2012)

6. Ding, L., Ley, T.J., Larson, D.E., Miller, C.A., Koboldt, D.C., Welch, J.S., Ritchey, J.K., Young, M.A., Lamprecht, T., McLellan, M.D., McMichael, J.F., Wallis, J.W., Lu, C., Shen, D., Harris, C.C., Dooling, D.J., Fulton, R.S., Fulton, L.L., Chen, K., Schmidt, H., Kalicki-Veizer, J., Magrini, V.J., Cook, L., McGrath, S.D., Vickery, T.L., Wendl, M.C., Heath, S., Watson, M.A., Link, D.C., Tomasson, M.H., Shannon, W.D., Payton, J.E., Kulkarni, S., Westervelt, P., Walter, M.J., Graubert, T.A., Mardis, E.R., Wilson, R.K., DiPersio, J.F.: Clonal evolution in relapsed acute myeloid leukaemia revealed by whole-genome sequencing. Nature 481(7382), 506–510 (2012)

7. Gillies, R.J., Verduzco, D., Gatenby, R.A.: Evolutionary dynamics of carcinogenesis and why targeted therapy does not work. Nat Rev Cancer 12(7), 487–493 (2012)

8. Burrell, R.A., McGranahan, N., Bartek, J., Swanton, C.: The causes and consequences of genetic heterogeneity in cancer evolution. Nature 501(7467), 338–345 (2013)

9. Gerstung, M., Beisel, C., Rechsteiner, M., Wild, P., Schraml, P., Moch, H., Beerenwinkel, N.: Reliable detection of subclonal single-nucleotide variants in tumour cell populations. Nature Communications 3 (2012)

10. Roth, A., Khattra, J., Yap, D., Wan, A., Laks, E., Biele, J., Ha, G., Aparicio, S., Bouchard-Cote, A., Shah, S.P.: PyClone: statistical inference of clonal population structure in cancer. Nat Meth 11(4), 396–398 (2014)

11. Ha, G., Roth, A., Khattra, J., Ho, J., Yap, D., Prentice, L.M., Melnyk, N., McPherson, A., Bashashati, A., Laks, E., Biele, J., Ding, J., Le, A., Rosner, J., Shumansky, K., Marra, M.A., Gilks, C.B., Huntsman, D.G., McAlpine, J.N., Aparicio, S., Shah, S.P.: TITAN: inference of copy number architectures in clonal cell populations from tumor whole-genome sequence data. Genome Research 24(11), 1881–1893 (2014)

12. Zare, H., Wang, J., Hu, A., Weber, K., Smith, J., Nickerson, D., Song, C., Witten, D., Blau, C.A., Noble, W.S.: Inferring clonal composition from multiple sections of a breast cancer. PLoS Comput Biol 10(7), 1–15 (2014)

13. El-Kebir, M., Oesper, L., Acheson-Field, H., Raphael, B.J.: Reconstruction of clonal trees and tumor composition from multi-sample sequencing data. Bioinformatics 31(12), 62–70 (2015)

14. Navin, N.: Cancer genomics: one cell at a time. Genome Biology 15(8), 452–465 (2014)

15. Jiao, W., Vembu, S., Deshwar, A.G., Stein, L., Morris, Q.: Inferring clonal evolution of tumors from single nucleotide somatic mutations. BMC Bioinformatics 15(1), 1–16 (2014)

16. Deshwar, A.G., Vembu, S., Yung, C.K., Jang, G.H., Stein, L., Morris, Q.: Phylowgs: Reconstructing subclonal composition and evolution from whole-genome sequencing of tumors. Genome Biology 16(1), 1–20 (2015)

17. El-Kebir, M., Satas, G., Oesper, L., Raphael, B.: Inferring the mutational history of a tumor using multi-state perfect phylogeny mixtures. Cell Systems 3(1), 43–53 (2016)

18. Jiang, Y., Qiu, Y., Minn, A.J., Zhang, N.R.: Assessing intratumor heterogeneity and tracking longitudinal and spatial clonal evolutionary history by next-generation sequencing. Proceedings of the National Academy of Sciences 113(37), 5528–5537 (2016)

19. Gerlinger, M., Rowan, A.J., Horswell, S., Larkin, J., Endesfelder, D., Gronroos, E., Martinez, P., Matthews, N., Stewart, A., Tarpey, P., Varela, I., Phillimore, B., Begum, S., McDonald, N.Q., Butler, A., Jones, D., Raine, K., Latimer, C., Santos, C.R., Nohadani, M., Eklund, A.C., Spencer-Dene, B., Clark, G., Pickering, L., Stamp, G., Gore, M., Szallasi, Z., Downward, J., Futreal, P.A., Swanton, C.: Intratumor heterogeneity and branched evolution revealed by multiregion sequencing. New England Journal of Medicine 366(10), 883–892 (2012)

20. Yates, L.R., Gerstung, M., Knappskog, S., Desmedt, C., Gundem, G., Van Loo, P., Aas, T., Alexandrov, L.B., Larsimont, D., Davies, H., Li, Y., Ju, Y.S., Ramakrishna, M., Haugland, H.K., Lilleng, P.K., Nik-Zainal, S., McLaren, S., Butler, A., Martin, S., Glodzik, D., Menzies, A., Raine, K., Hinton, J., Jones, D., Mudie, L.J., Jiang, B., Vincent, D., Greene-Colozzi, A., Adnet, P.-Y., Fatima, A., Maetens, M., Ignatiadis, M., Stratton, M.R., Sotiriou, C., Richardson, A.L., Lonning, P.E., Wedge, D.C., Campbell, P.J.: Subclonal diversification of primary breast cancer revealed by multiregion sequencing. Nat Med 21(7), 751–759 (2015). Article

21. Navin, N.E.: The first five years of single-cell cancer genomics and beyond. Genome Res 25(10), 1499–1507 (2015)

22. Hou, Y., Song, L., Zhu, P., Zhang, B., Tao, Y., Xu, X., Li, F., Wu, K., Liang, J., Shao, D., Wu, H., Ye, X., Ye, C., Wu, R., Jian, M., Chen, Y., Xie, W., Zhang, R., Chen, L., Liu, X., Yao, X., Zheng, H., Yu, C., Li, Q., Gong, Z., Mao, M., Yang, X., Yang, L., Li, J., Wang, W., Lu, Z., Gu, N., Laurie, G., Bolund, L., Kristiansen, K., Wang, J., Yang, H., Li, Y., Zhang, X., Wang, J.: Single-cell exome sequencing and monoclonal evolution of a JAK2-negative myeloproliferative neoplasm. Cell 148(5), 873–885 (2012)

23. Xu, X., Hou, Y., Yin, X., Bao, L., Tang, A., Song, L., Li, F., Tsang, S., Wu, K., Wu, H., He, W., Zeng, L., Xing, M., Wu, R., Jiang, H., Liu, X., Cao, D., Guo, G., Hu, X., Gui, Y., Li, Z., Xie, W., Sun, X., Shi, M., Cai, Z., Wang, B., Zhong, M., Li, J., Lu, Z., Gu, N., Zhang, X., Goodman, L., Bolund, L., Wang, J., Yang, H., Kristiansen, K., Dean, M., Li, Y., Wang, J.: Single-cell exome sequencing reveals single-nucleotide mutation characteristics of a kidney tumor. Cell 148(5), 886–895 (2012)

24. Wang, Y., Waters, J., Leung, M.L., Unruh, A., Roh, W., Shi, X., Chen, K., Scheet, P., Vattathil, S., Liang, H., Multani, A., Zhang, H., Zhao, R., Michor, F., Meric-Bernstam, F., Navin, N.E.: Clonal evolution in breast cancer revealed by single nucleus genome sequencing. Nature 512(7513), 155–160 (2014)

25. Gawad, C., Koh, W., Quake, S.R.: Dissecting the clonal origins of childhood acute lymphoblastic leukemia by single-cell genomics. Proceedings of the National Academy of Sciences 111(50), 17947–17952 (2014)

26. Li, Y., Xu, X., Song, L., Hou, Y., Li, Z., Tsang, S., Li, F., Im, K., Wu, K., Wu, H., Ye, X., Li, G., Wang, L., Zhang, B., Liang, J., Xie, W., Wu, R., Jiang, H., Liu, X., Yu, C., Zheng, H., Jian, M., Nie, L., Wan, L., Shi, M., Sun, X., Tang, A., Guo, G., Gui, Y., Cai, Z., Li, J., Wang, W., Lu, Z., Zhang, X., Bolund, L., Kristiansen, K., Wang, J., Yang, H., Dean, M., Wang, J.: Single-cell sequencing analysis characterizes common and cell-lineage-specific mutations in a muscle-invasive bladder cancer. GigaScience 1(1), 12 (2012)

27. Zafar, H., Wang, Y., Nakhleh, L., Navin, N., Chen, K.: Monovar: single-nucleotide variant detection in single cells. Nat Meth 13(6), 505–507 (2016)

28. Zhang, C.-Z., Adalsteinsson, V.A., Francis, J., Cornils, H., Jung, J., Maire, C., Ligon, K.L., Meyerson, M., Love, J.C.: Calibrating genomic and allelic coverage bias in single-cell sequencing. Nature Communications 6, 6822 (2015)

29. Navin, N., Kendall, J., Troge, J., Andrews, P., Rodgers, L., McIndoo, J., Cook, K., Stepansky, A., Levy, D., Esposito, D., Muthuswamy, L., Krasnitz, A., McCombie, W.R., Hicks, J., Wigler, M.: Tumour evolution inferred by single-cell sequencing. Nature 472(7341), 90–94 (2011)

30. Baslan, T., Kendall, J., Rodgers, L., Cox, H., Riggs, M., Stepansky, A., Troge, J., Ravi, K., Esposito, D., Lakshmi, B., Wigler, M., Navin, N., Hicks, J.: Genome-wide copy number analysis of single cells. Nat. Protocols 7(6), 1024–1041 (2012)

31. Leung, M.L., Wang, Y., Kim, C., Gao, R., Jiang, J., Sei, E., Navin, N.E.: Highly multiplexed targeted dna sequencing from single nuclei. Nat. Protocols 11(2), 214–235 (2016). Protocol

32. Macosko, E., Basu, A., Satija, R., Nemesh, J., Shekhar, K., Goldman, M., Tirosh, I., Bialas, A., Kamitaki, N., Martersteck, E., Trombetta, J., Weitz, D., Sanes, J., Shalek, A., Regev, A., McCarroll, S.: Highly parallel genome-wide expression profiling of individual cells using nanoliter droplets. Cell 161(5), 1202–1214 (2015)

33. Yu, C., Yu, J., Yao, X., Wu, W.K., Lu, Y., Tang, S., Li, X., Bao, L., Li, X., Hou, Y., Wu, R., Jian, M., Chen, R., Zhang, F., Xu, L., Fan, F., He, J., Liang, Q., Wang, H., Hu, X., He, M., Zhang, X., Zheng, H., Li, Q., Wu, H., Chen, Y., Yang, X., Zhu, S., Xu, X., Yang, H., Wang, J., Zhang, X., Sung, J.J., Li, Y., Wang, J.: Discovery of biclonal origin and a novel oncogene slc12a5 in colon cancer by single-cell sequencing. Cell Res 24(6), 701–712 (2014)

34. Eirew, P., Steif, A., Khattra, J., Ha, G., Yap, D., Farahani, H., Gelmon, K., Chia, S., Mar, C., Wan, A., Laks, E., Biele, J., Shumansky, K., Rosner, J., McPherson, A., Nielsen, C., Roth, A.J.L., Lefebvre, C., Bashashati, A., de Souza, C., Siu, C., Aniba, R., Brimhall, J., Oloumi, A., Osako, T., Bruna, A., Sandoval, J.L., Algara, T., Greenwood, W., Leung, K., Cheng, H., Xue, H., Wang, Y., Lin, D., Mungall, A.J., Moore, R., Zhao, Y., Lorette, J., Nguyen, L., Huntsman, D., Eaves, C.J., Hansen, C., Marra, M.A., Caldas, C., Shah, S.P., Aparicio, S.: Dynamics of genomic clones in breast cancer patient xenografts at single-cell resolution. Nature 518(7539), 422–426 (2015)

35. Huelsenbeck, J.P., Ronquist, F.: Mrbayes: Bayesian inference of phylogenetic trees. Bioinformatics 17(8), 754–755 (2001)

36. Yuan, K., Sakoparnig, T., Markowetz, F., Beerenwinkel, N.: Bitphylogeny: a probabilistic framework for reconstructing intra-tumor phylogenies. Genome Biology 16(1), 1–16 (2015)

37. Ross, E.M., Markowetz, F.: OncoNEM: inferring tumor evolution from single-cell sequencing data. Genome Biology 17(1), 1–14 (2016)

38. Jahn, K., Kuipers, J., Beerenwinkel, N.: Tree inference for single-cell data. Genome Biology 17(1), 1–17 (2016)

39. Kim, K.I., Simon, R.: Using single cell sequencing data to model the evolutionary history of a tumor. BMC Bioinformatics 15(1), 27 (2014)

40. Ma, J., Ratan, A., Raney, B.J., Suh, B.B., Miller, W., Haussler, D.: The infinite sites model of genome evolution. Proceedings of the National Academy of Sciences 105(38), 14254–14261 (2008)

41. Gusfield, D.: Algorithms on Strings, Trees and Sequences: Computer Science and Computational Biology. Cambridge University Press, Cambridge (1997)

42. Davis, A., Navin, N.E.: Computing tumor trees from single cells. Genome Biology 17(1), 1–4 (2016)

43. Yang, Z., Rannala, B.: Molecular phylogenetics: principles and practice. Nat Rev Genet 13(5), 303–314 (2012)

44. DePristo, M.A., Banks, E., Poplin, R., Garimella, K.V., Maguire, J.R., Hartl, C., Philippakis, A.A., del Angel, G., Rivas, M.A., Hanna, M., McKenna, A., Fennell, T.J., Kernytsky, A.M., Sivachenko, A.Y., Cibulskis, K., Gabriel, S.B., Altshuler, D., Daly, M.J.: A framework for variation discovery and genotyping using next-generation DNA sequencing data. Nat Genet 43(5), 491–498 (2011)

45. Robinson, D.F., Foulds, L.R.: Comparison of phylogenetic trees. Mathematical Biosciences 53(1), 131–147 (1981)

46. Wu, H., Zhang, X.-Y., Hu, Z., Hou, Q., Zhang, H., Li, Y., Li, S., Yue, J., Jiang, Z., Weissman, S.M., Pan, X., Ju, B.-G., Wu, S.: Evolution and heterogeneity of non-hereditary colorectal cancer revealed by single-cell exome sequencing. Oncogene (2016)

47. Gusfield, D.: ReCombinatorics: The Algorithmics of Ancestral Recombination Graphs and Explicit Phylogenetic Networks. The MIT Press, Cambridge (2014)

48. Pupko, T., Pe, I., Shamir, R., Graur, D.: A fast algorithm for joint reconstruction of ancestral amino acid sequences. Molecular Biology and Evolution 17(6), 890–896 (2000)

49. Leung, M.L., Davis, A., Gao, R., Casasent, A., Wang, Y., Sei, E., Sanchez, E., Maru, D., Kopetz, S., Navin, N.E.: Single cell dna sequencing reveals a late-dissemination model in metastatic colorectal cancer. Genome Research (2017)

50. Gao, R., Davis, A., McDonald, T.O., Sei, E., Shi, X., Wang, Y., Tsai, P.-C., Casasent, A., Waters, J., Zhang, H., Meric-Bernstam, F., Michor, F., Navin, N.E.: Punctuated copy number evolution and clonal stasis in triple-negative breast cancer. Nat Genet 48(10), 1119–1130 (2016)

51. Baslan, T., Kendall, J., Ward, B., Cox, H., Leotta, A., Rodgers, L., Riggs, M., D’Italia, S., Sun, G., Yong, M., Miskimen, K., Gilmore, H., Saborowski, M., Dimitrova, N., Krasnitz, A., Harris, L., Wigler, M., Hicks, J.: Optimizing sparse sequencing of single cells for highly multiplex copy number profiling. Genome Research 25(5), 714–724 (2015)

52. Navin, N., Hicks, J.: Future medical applications of single-cell sequencing in cancer. Genome Medicine 3(5), 31 (2011)

53. Wang, Y., Navin, N.: Advances and applications of single-cell sequencing technologies. Molecular Cell 58(4), 598–609 (2015)

54. Felsenstein, J.: Evolutionary trees from dna sequences: A maximum likelihood approach. Journal of Molecular Evolution 17(6), 368–376 (1981)

55. Hastings, W.K.: Monte carlo sampling methods using markov chains and their applications. Biometrika 57(1), 97–109 (1970)

56. Lakner, C., van der Mark, P., Huelsenbeck, J.P., Larget, B., Ronquist, F.: Efficiency of markov chain monte carlo tree proposals in bayesian phylogenetics. Systematic Biology 57(1), 86–103 (2008)

57. Than, C., Ruths, D., Nakhleh, L.: PhyloNet: a software package for analyzing and reconstructing reticulate evolutionary relationships. BMC Bioinformatics 9(1), 322 (2008). DOI:10.1186/1471-2105-9-322.

58. Imam, S., Sarkar, V.: Habanero-java library: A java 8 framework for multicore programming. In: Proceedings of the 2014 International Conference on Principles and Practices of Programming on the Java Platform: Virtual Machines, Languages, and Tools. PPPJ’14, pp. 75–86. ACM, New York, NY, USA (2014)

